# A Germinal Center–Independent Innate-Like Memory B Cell Compartment of B1 Origin

**DOI:** 10.64898/2026.07.22.740083

**Authors:** Paula M. Villavicencio, Maria Bottermann, Myriam Ortiz Isuiza, Shivang Sunil Parikh, John E. Warner, Anthony Alicea, Elizabeth Zhou, Thavaleak Prum, Hannah Naili, Xiaotie Liu, Stephanie R. Weldon, Facundo D. Batista

## Abstract

Memory B cells (MBCs) are a critical cellular reservoir for long-term humoral immunity. MBCs display heterogeneous isotypes and surface markers and can arise through both germinal center (GC)-dependent and GC-independent extrafollicular (EF) pathways. Both the mechanisms controlling EF MBC differentiation and the identities of the B cell populations from which EF MBCs derive remain poorly understood. To capture MBC diversity, we applied a broad selection strategy followed by transcriptional profiling, identifying a subset of MBCs characterized by minimal class switching, limited somatic hypermutation, and an innate-like gene signature. Using genetic models, we demonstrated that this subset arises independently of GC responses and derives from innate B1 cells. These innate-like MBCs differentiate into antigen-specific antibody-secreting cells and confer protection against lethal viral infection. Together, our findings define a previously unrecognized arm of MBC responses, demonstrating that innate B1 cells contribute a non-redundant antibody repertoire to protective immunological memory.

## INTRODUCTION

One of the main goals of modern vaccinology is the induction of potent immune memory. The capacity to elicit long-term protection after pathogen encounter is essential to prevent recurring infections and establish population-level herd immunity. Such durable long-term protection can be achieved by the generation of memory B cell (MBC) responses; thus, an in-depth understanding of MBC development and heterogeneity is highly relevant in the context of vaccination.

Upon antigen encounter, B cells interact with T cells extrafollicularly. Immediately after this initial activation, B cells follow diverse fates, including migration into the follicle to form germinal centers (GCs), early plasmablast differentiation, or emergence as a first wave of MBCs prior to GC formation (Pape et al. 2011; Taylor, Pape, and Jenkins 2012; Viant et al. 2021; F. J. Weisel et al. 2016). These early MBCs are mostly unmutated and non-class-switched (Kaji et al. 2012; Taylor, Pape, and Jenkins 2012; F. J. Weisel et al. 2016). To identify murine MBCs by flow cytometry, studies have classically relied on the selection of class-switched or fate-mapped cells (Haase et al. 2022; Mesin et al. 2020), which may exclude this early wave of MBCs. Alternatively, surface markers such as PDL2 and CD80 have been used (Tomayko et al. 2010; F. Weisel and Shlomchik 2017; N. M. Weisel et al. 2022; Zuccarino-Catania et al. 2014), which may not capture all MBC subtypes or resolve their heterogeneity (Callahan et al. 2024).

More recent studies using single-cell sequencing have revealed that MBCs are more diverse and complex than previously appreciated (Cooper et al. 2024; Gregoire et al. 2022; Riedel et al. 2020; Risley et al. 2025). Among the subsets studied in detail in recent years are CD11c⁺T-bet⁺ MBCs, which were originally identified in response to viral infection and have since been extensively characterized using transcriptomic and functional approaches (Johnson et al. 2020; Risley et al. 2021, 2025; Rubtsova et al. 2013; Song et al. 2022; Stone et al. 2019). In addition, newly described MBC subsets include populations defined by an interferon-stimulated gene (ISG) signature in the context of chronic viral infection (Cooper et al. 2024), as well as CCR6^+^ bona fide antigen-specific and bystander memory populations distinguished by CXCR3-expression (Gregoire et al. 2022). However, many recent studies selected MBCs based on activation-induced cytidine deaminase (AID) expression, class switching, antigen specificity, or, when starting from broader populations, the use of restricted analysis potentially excluding additional MBC subsets.

To achieve a comprehensive view of murine MBCs, we employed a multi-strategy approach to enrich MBCs regardless of their class-switch status, mutation load, or antigen-specificity. Utilizing single-cell sequencing and fate-mapping, we identified an MBC population with a unique transcriptional profile, distinguished by CD11b and CD24 expression. These MBCs exhibited low levels of somatic hypermutation and class-switching, reminiscent of early GC-independent MBCs. Using complementary mouse models, we demonstrated that these cells arise via an extrafollicular path. Their absence in CD19 KO mice suggested a B1 innate origin, which was supported by shared VH/VL pairs between B1 cells and innate-like CD11b/CD24-expressing MBCs, and further confirmed through congenic adoptive transfer experiments. Finally, we demonstrated that these B1-derived innate-like MBCs efficiently differentiated into antigen-specific antibody secreting cells (ASCs) and were able to protect from lethal viral infection *in vivo*. Thus, innate-like MBCs provide a protective, non-redundant antibody repertoire during recall responses.

## RESULTS

### A broad selection strategy captures memory B cell gene expression diversity

To define the diversity of memory B cells (MBCs), we applied multiple complementary enrichment strategies in wild-type and AID-reporter mice (referred to as AID-tdTomato mice) (Madisen et al. 2010; Robbiani et al. 2008) following NP-CGG immunization (**Fig. 1A**). For initial MBC selection, we deployed previously described MBC markers: B220^+^CD138^−^CD95^int^CD38^+^IgD^−^ (Ridderstad and Tarlinton 1998; Aiba et al. 2010; Talay et al. 2013). Downstream of this general selection approach, we specifically sorted cells using four different surface protein expression schemes: 1) PDL2^+^, 2) CD38^+^NP^+^, 3) AID-tdTomato^+^ and 4) CD38^+^NP^+^IgM^+^ and CD38^+^NP^+^IgG^+^ (Tomayko et al. 2010; Zuccarino-Catania et al. 2014). These populations were sorted across 6 independent experiments. For each set, we performed scRNA-seq and then integrated the 6 datasets using Harmony (Korsunsky et al. 2019) (**Fig. 1B**). Clusters were generated using the Louvain algorithm and clusters containing unrelated cell types, such as non-B cells, plasma cells (PC), germinal centers (GC), naïve and transitional B cells (**Fig. S1A–D**) were removed, resulting in 9128 retained MBCs. We identified seven MBC clusters (**Fig. 1C**), each defined by distinctive differentially expressed genes (DEGs) (**Fig. 1D**), that allowed us to classify them into five major cell types distinguished by divergent transcriptional programs. Firstly, the broadly studied classical MBCs, which are characterized by CD23, CCR6, *Bach2* and *Hhex* expression (Duan et al. 2021; Elgueta et al. 2015; Laidlaw et al. 2020; Shinnakasu et al. 2016; Suan et al. 2017), were distributed among three clusters in our dataset: 1, 5 and 7, all of which expressed high levels of *Ccr6*, *Hhex* and *Fcer2a* (**Fig. 1C–D, S1E–F**). MBCs expressing marginal zone (MZ)-like features, such as *Cr2, Cd1d1*, and *S1pr3* (Amano et al. 1998; Hendricks, Bos, and Kroese 2018; Takahashi et al. 1997; Tedford et al. 2017) corresponded to cluster 2. Cluster 3 presented innate-like gene expression, such as *S100a6 and Nfam1* (Juchem et al. 2021; Kozlyuk et al. 2019; Tong et al. 2023), alongside the MBC-associated transcriptional repressor *Zbtb32* (Jash et al. 2016, 2019). Cluster 4 was characterized by the expression of *Apoe, Fcer1g,* and atypical B cell associated gene *Zeb2 (Dai et al. 2024; Gao et al. 2024).* Despite its low *Tbx21* and *Itgax* expression, other genes associated with Tbet^+^CD11c^+^ MBCs (Song et al. 2022) were enriched (**Fig. S1E**). Lastly, cluster 6 displayed increased expression of *Nfkbid, Il4I1 and Ncl (***Fig. 1C–D**), which are involved in modulation of antibody responses and isotype class switching, B cell activation, and ribosome biogenesis, respectively (Arnold et al. 2012; Bod et al. 2018; Fremerey et al. 2016; Hanakahi et al. 1997; Khoenkhoen et al. 2022; Souza et al. 2021). Notably, we were also able to identify an ISG^+^ MBC subset (Cooper et al. 2024) in select experiments, albeit in very small numbers (**Fig. S2A–C).**

**Figure 1.**
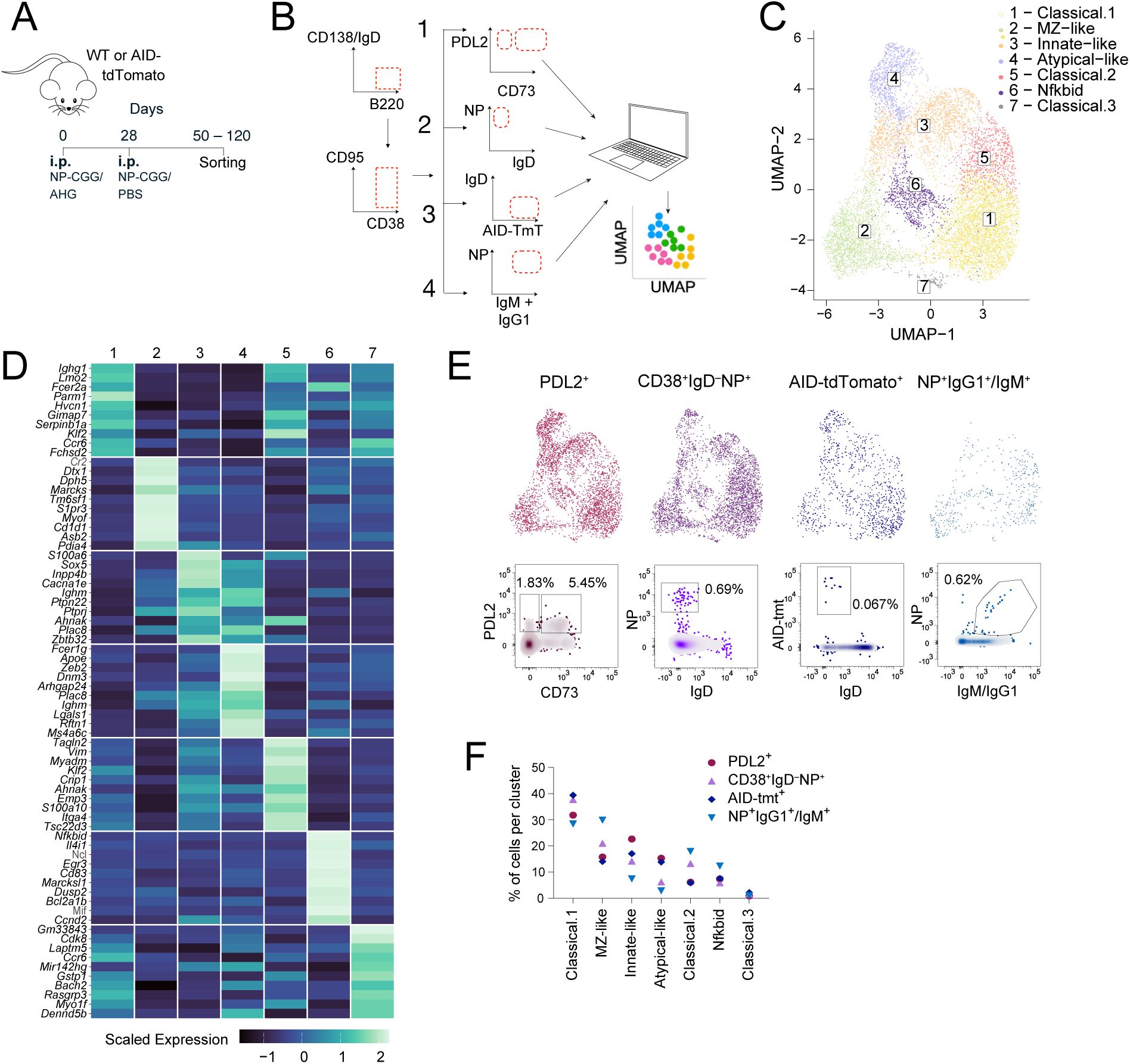
Broad selection of memory B cells from spleen for scRNAseq analysis. (A) Schematic of immunization and boost model. WT or AID-tdTomato mice were immunized (day 0) intraperitoneally (i.p.) with 50 µg NP-CGG in alhydrogel (AHG) and boosted (day 28) with 50 µg NP-CGG in PBS. Memory B cells (MBCs) were isolated on days 50–120 and enriched using strategies described in (B). (B) Schematic of four MBC isolation strategies used. All cells were first gated as: B220^+^CD138^−^CD95^int^CD38^+^IgD^−^. Cells then underwent one of four sorting strategies: 1) PDL2^+^: sorted as total PDL2^+^IgD^−^, PDL2^+^CD73^+^IgD^−^ or PDL2^+^CD73^−^IgD^−^; 2) Antigen specific: CD38^+^IgD^−^NP^+^; 3) Class switched: AID-tdTomato^+^IgD^−^; 4) Antigen specific 2: NP^+^IgM^+^ and NP^+^IgG1^+^. Individual replicates sorted per condition across 6 independent experiments: 1) 7 samples, 2) 16 samples, 3) 3 samples, 4) 3 samples. During single-cell analysis using Seurat, all datasets were integrated using Harmony. (C) UMAP showing Louvain clustering of MBCs sorted as in (B) at resolution 0.5, following Harmony integration of the six datasets and removal of non-B cells, plasma cells, germinal center, naïve, and transitional B cells. (D) Heatmap showing the top 10 gene expression markers for each cluster. Color key: scaled mean expression. (E) *Top*: UMAP showing MBCs faceted by sorting condition described in (B). *Bottom*: flow cytometry plots representing gating strategies shown in (B) used to isolate MBCs displayed in the UMAPs above. (F) Quantification of MBC frequencies in each cluster by isolation method.

Stratifying the UMAP by sorting condition revealed a homogenous distribution of MBCs along the 7 clusters for each of the gating strategies used (**Fig. 1E**). No significant difference was found in the frequency of cells per cluster between sorting conditions (**Fig. 1F**), indicating that we obtained a robust representation of MBC clusters throughout the sorting strategies. Subsequently, MBCs were selected using the CD38^+^, PDL2^+^, and IgD^−^gating scheme, which allows us to obtain MBCs regardless of their antigen specificity, affinity maturation or class switching state.

Together, these data define a robust and heterogeneous landscape of MBCs, revealing multiple transcriptionally distinct subsets that are consistently captured across independent enrichment strategies.

### BCR variety across transcriptionally defined MBC populations

To further investigate lineage diversity, we selected four MBC populations: classical, MZ-like, innate-like and atypical-like MBCs, and performed single-cell BCR-seq analysis on the previously integrated dataset (**Fig. 1**). We first evaluated variable gene (V gene) usage across these four populations (3569 MBCs in total), focusing on the 10 most frequent V genes. Classical and innate-like MBCs showed a more restricted V gene repertoire, with three variable heavy chain (VH) genes accounting for ∼47% and ∼35% of classical and innate-like MBCs, respectively. Similarly, variable light chains (VL) were dominated by three V genes in ∼57% and ∼44% of these respective MBCs. In contrast, MZ-like and atypical-like MBCs exhibited a broader V gene repertoire (**Fig. 2A**). Given these differences in V gene repertoires among MBC populations, we sought to examine their mutational load across both VH and VL genes. We found distinct ranges of amino acid (aa) changes across the different MBC populations. The classical MBC population showed the highest number of aa changes in the VH (mean of 4.5) and VL (mean of 1.7) chains. In contrast, MZ-like, innate-like, and atypical-like MBCs contained fewer aa changes (VH means of 0.97, 1.50, and 1.55, VL means of 0.44, 0.71, and 0.74, respectively) (**Fig. 2B**).

**Figure 2.**
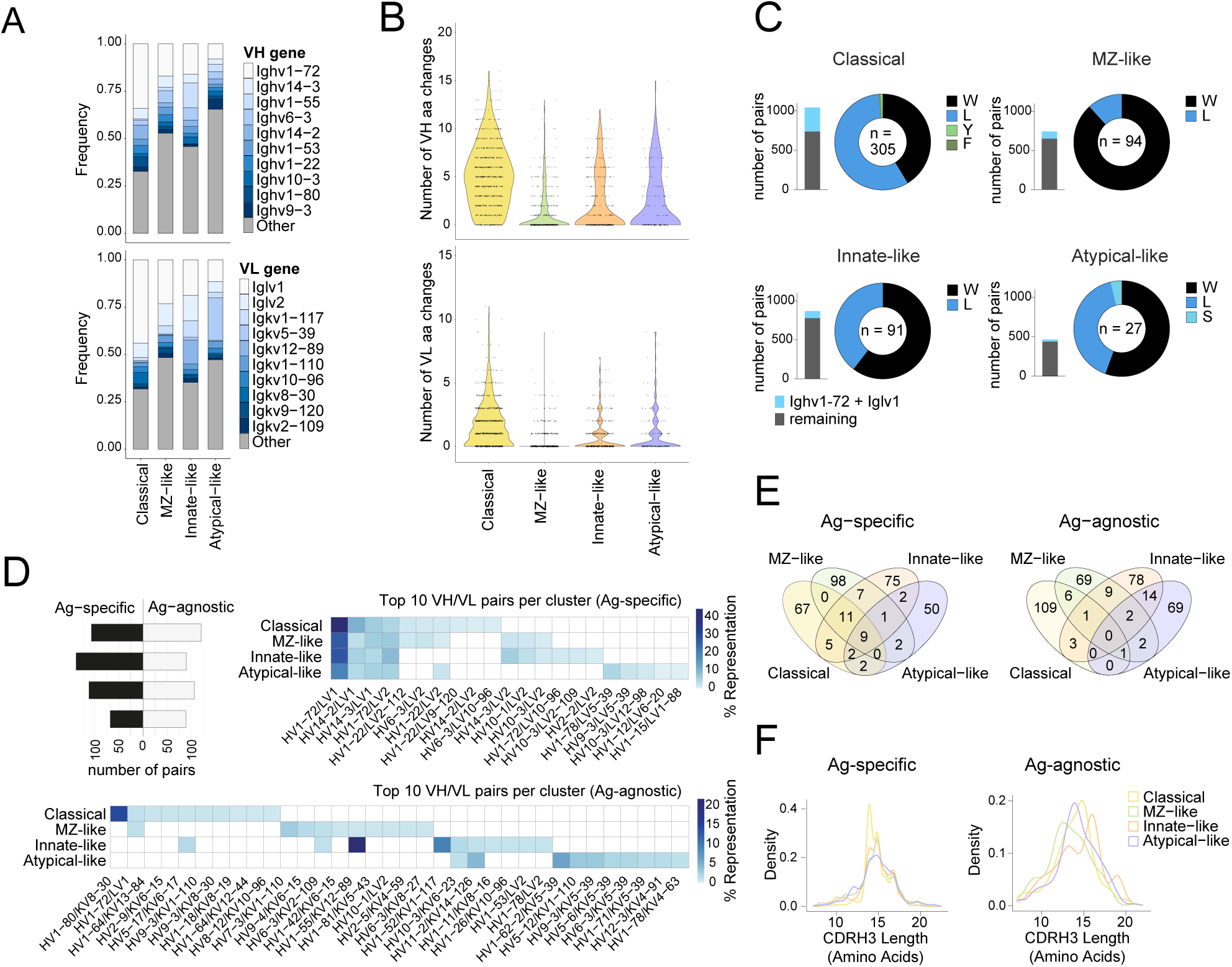
BCR diversity across four selected MBC populations. (A) Top 10 VH (top) and VL (bottom) genes across classical (1207 cells), MZ-like (914 cells), innate-like (947 cells), and atypical-like (501 cells) MBC clusters. (B) Violin plots showing number of amino acid (aa) changes in variable heavy (VH) chain (*top*) and variable light (VL) chain (*bottom*) per indicated cluster. (C) Bar plots quantifying the number of Ighv1-72/Iglv1 pairs and remaining VH/VL pairs found in the four selected MBC clusters. Donut charts showing the amino acid composition at position 33 of Ighv1-72. n = total number of Ighv1-72/Iglv1 pairs analyzed per cluster. (D) *Top left*: Number of VH/VL pairs per indicated cluster, split by antigen-specific (black) and antigen-agnostic (white) cells. Top 10 VH/VL pairs per indicated cluster in antigen-specific (*top right*) and antigen-agnostic (*bottom*) cells per indicated cluster. (E) Venn diagram showing overlap of VH/VL pairs between the indicated MBCs in antigen-specific (*left*) and antigen-agnostic (*right)* cells. (F) CDRH3 length distribution in amino acids of the indicated MBC populations in antigen-specific (*left*) and antigen-agnostic (*right*) cells.

To determine whether the number of mutations in VH and VL chains correlated with affinity maturation, we analyzed the Ighv1-72 / Iglv1 canonical NP-reactive germline pair, which were the most frequently used VH and VL genes across all populations (**Fig. 2A**). We quantified the representation of this VH/VL pair in the four MBC types, as well as the frequency of the W33L mutation in the CDRH1, which is known to increase affinity for NP by 10-fold (Allen et al. 1988; Cumano and Rajewsky 1986). Classical MBCs contained the highest number of Ighv1-72 / Iglv1 VH/VL pairs (∼29%) with approximately 57% of those containing the W33L mutation, demonstrating that this subset largely consists of affinity-matured NP-binders (**Fig. 2C**). In contrast, the Ighv1-72 / Iglv1 pair comprised only 6–12% of MZ-like, innate-like, and atypical-like MBC VH/VL pairs. Approximately 40% of Ighv1-72 contained the W33L high affinity mutation in innate and atypical-like MBCs, while MZ-like MBCs had only ∼12% (**Fig. 2C**). Therefore, MZ, innate-like, and atypical-like MBCs exhibit reduced affinity to NP as assessed by W33L mutation frequency.

To further interrogate BCR diversity of these four MBC populations, we evaluated whether antigen-specificity affected isotype switching and VH/VL pairing. Classical MBCs showed increased class-switching compared to the remaining subsets (**Fig. S2D**), which is consistent with its increased mutation load given that both processes rely on AID (Leeman-Neill, Lim, and Basu 2018). Antigen-specific MBCs are overall biased towards IgG1, coupled with an increased use of lambda light chains (**Fig. S2D, left**), the latter being a distinguishing feature of NP-binders (Reth, Hämmerling, and Rajewsky 1978). In contrast, antigen-agnostic MBCs remained mostly unswitched, except for classical MBCs, which exhibited near-complete class switching. Antigen-agnostic MBCs also showed a more typical Igk/Igl distribution, with only MZ-like and innate-like MBCs exhibiting elevated Igl usage over the canonical ∼5% (**Fig. S2D, right**). VH/VL pairing analysis showed that antigen-specific MBCs, regardless of their subset, were dominated by NP-binding VH/VL pairs, while antigen-agnostic MBCs showed increased pairing diversity. Antigen-agnostic innate-like MBCs retained a dominant use of two VH/VL pairs observed in other antigen-agnostic MBCs: Ighv1-55 / Igkv12-89 and Ighv1-52 / Igkv1-117 (**Fig. 2D**). Congruent with the VH/VL pairing diversity observed, a total of nine VH/VL pairs were shared among all antigen-specific MBCs, while reduced sharing was found between antigen-agnostic populations (**Fig. 2E**). Additionally, CDRH3 length analysis of antigen-specific MBCs revealed three distinct peaks across the four populations (**Fig. 2F, left**), corresponding to the CDRH3 length associated with the three dominant VH genes (**Fig. S2E**). Conversely, antigen-agnostic MBCs exhibited a monophasic length distribution, except for innate-like MBCs which showed three peaks (**Fig. 2F, right**) corresponding to the CDRH3 length of their preferred VH genes (**Fig. S2F**), underlining a biased VH/VL pair usage.

In summary, our broad approach in enriching MBCs allowed us to consistently identify four major populations that differed in their transcriptional and mutational status, as well as to establish an enrichment method that avoids bias towards class-switching, somatic hypermutation or restricted antigen binding. Notably, among these discrete populations, we isolated an innate-like MBC population that, although detected in other sc-RNAseq datasets (Gregoire et al. 2022; Riedel et al. 2020; Risley et al. 2025), has not yet been studied.

### Innate-like MBCs develop independent of GCs

The increased mutational load, affinity maturation and class switching found in classical MBCs is suggestive of a GC-dependent origin, while the reduced somatic hypermutation and class switching observed in the innate-like MBC population, as well as MZ-like and atypical-like MBCs, indicates that these three subsets may arise GC-independently. To interrogate this hypothesis, we took advantage of two complementary mouse models: one incapable of forming GCs, and a second where GC B cells are irreversibly marked.

The first model, Bcl6^fl/fl^ CD4 Cre^+/–^, has a CD4 T cell-specific Bcl6 deficiency and therefore lacks T follicular helper cells. These mice should be unable to form GCs, while GC-independent B cell activation should remain intact. Consequently, to obtain MBCs of GC-independent origin, we generated mixed bone marrow (BM) chimeras by reconstituting BM-depleted RAG1-KO mice with a mix of RAG1-KO BM and either Bcl6^fl/fl^ CD4 Cre^−/–^ (Bcl6 CD4 WT) or Bcl6^fl/fl^ CD4 Cre^+/–^ (Bcl6 CD4 KO) BM (**Fig. 3A**). We confirmed Tfh cell deficiency in Bcl6 CD4 KO chimeras (**Fig. S3A**) and observed the consequent GC impairment, while extrafollicular plasmablast differentiation remained intact (**Fig. S3B).**

**Figure 3.**
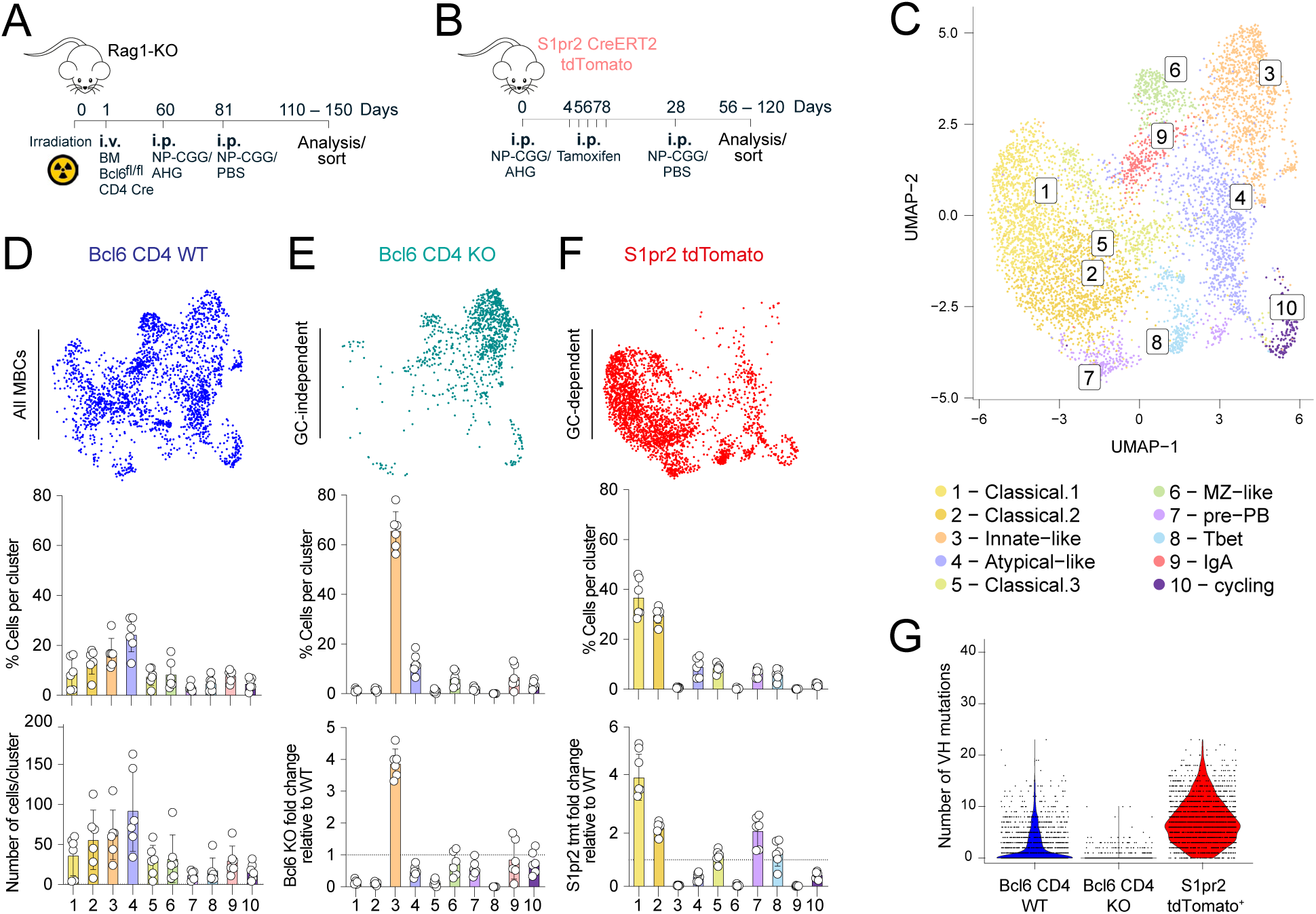
Identification of GC-dependent and GC-independent MBC populations. (A) Schematic of mouse model and immunization protocol used to evaluate GC-independent memory response. Chimeras were generated with a mix of 75% RAG1-KO and 25% Bcl6^fl/fl^ CD4 Cre^−/−^ (Bcl6 CD4 WT) or 25% Bcl6^fl/fl^ CD4 Cre^+/−^ (Bcl6 CD4 KO) bone marrow. For scRNA-seq, cells were isolated from Bcl6 CD4 WT mice (n = 7 mice) and Bcl6 CD4 KO mice (n = 10 mice) and processed as six biological replicates per model. (B) Schematic of mouse model and immunization protocol used to evaluate the GC-dependent memory response. S1pr2 Cre^ERT2^ tdTomato mice were injected i.p. with 2 mg of tamoxifen daily from days 4–8 post primary immunization. For scRNA-seq, MBCs from 12 S1pr2 Cre^ERT2^ tdTomato mice were pooled to generate six biological samples (two mice per sample). (A and B) Mice were immunized i.p. (day 0) with 50 µg NP-CGG/AHG, boosted (day 28) with 50 µg NP-CGG/PBS and sacrificed 4–8 weeks post-boost. (C) UMAP showing MBCs obtained from both GC-dependent and GC-independent models described in (A, B). Seurat clusters were obtained by Louvain clustering at resolution 0.5 after Harmony integration of the two datasets and removal of non-B cells, plasma cells, germinal center, naïve, and transitional B cells. (D–F) Top: UMAP split by mouse model showing cluster distribution of Bcl6 CD4 WT cells (D), Bcl6 CD4 KO cells (E), and S1pr2-tdTomato cells (F). Middle: Bar plots showing frequency of cells per cluster. Bottom: Bar plots showing number of cells per cluster (D) or the fold change in cluster-specific cell frequencies relative to WT Bcl6 CD4 WT cells (E, F). (G) Violin plots showing number of mutations in the VH chain of each mouse model.

The second model deployed, S1pr2 Cre^ERT2^ Rosa26-tdTomato strain (S1pr2 tdTomato), allowed us to determined which MBCs in our system were GC-derived through temporal tracing via a tamoxifen-inducible S1pr2-Cre expressed exclusively in GC B cells (Green et al. 2011; Shinnakasu et al. 2016) (**Fig. 3B**).

In both Bcl6 CD4 chimeras and S1pr2 tdTomato mice, we induced MBCs by using prime and boost NP-CGG immunization (**Fig. 3A–B**). S1pr2 mice were additionally treated with tamoxifen on days 4–8 after prime (**Fig. 3B**), irreversibly marking MBCs that were present in the GC at the time of tamoxifen treatment through tdTomato expression. MBCs from both mouse models were recovered approximately 100 days after primary immunization. MBCs from Bcl6 chimeras were sorted as NP^+^, CD38^+^, or PDL2^+^, while cells from the S1pr2 tomato model were sorted by tdTomato expression (**Fig. S3C**). Single-cell sequencing of sorted cells revealed similar MBC clusters to those observed in WT mice above (**Figs. 1C, 3C–D, S3D, S4A–B**), as well as additional clusters corresponding to pre-plasmablasts, Tbet^+^ MBCs, IgA^+^ MBCs, and cycling B cells (clusters 7 through 10, respectively).

We stratified the integrated UMAP by animal model and analyzed cluster distribution. Within WT control (Bcl6 CD4 WT) cells, we observed that ∼25% of cells were distributed in classical MBC clusters, while ∼8%, ∼16%, and 24% corresponded to MZ-like, innate-like, and atypical-like MBCs, respectively (**Fig. 3D**). In contrast, GC-independent Bcl6 CD4 KO-derived cells showed a minor fraction of classical MBCs (3.9%) but were predominantly associated with innate-like MBCs, accounting for 65% of cells, a ∼4-fold increase in frequency over WT mice. (**Fig. 3E**). MZ-like and atypical-like MBCs were also present among Bcl6 CD4 KO-derived cells, albeit at lower levels than in WT, representing 6% and 12% of cells respectively. Together, these data strongly support that MZ-like, atypical and mainly innate-like MBCs can develop independently of GC formation.

In the S1pr2 tdTomato^+^ GC-lineage traced cells, cluster analysis showed that 73% of GC-derived tdTomato^+^ cells mapped to classical MBC clusters. Consistently, there is a ∼4-to 2-fold increase in frequency of classical MBCs in S1pr2 tdTomato^+^ relative to WT controls (**Fig. 3F**). Atypical-like MBCs were present at lower frequencies (9%) than in WT (**Figs. 3D–F, S4C**), and MZ-like and innate-like MBCs were minimally represented (0.36% and 0.6%, respectively) (**Fig. 3F**). Together, these models indicate that MZ-like and particularly innate-like MBCs arose primarily through GC-independent pathways, whereas classical MBCs were largely GC-derived. The similar proportion of atypical-like MBCs found among Bcl6 CD4 KO-derived and tdTomato^+^ cells (12% and 9%, respectively) suggests a mixed origin for this population.

To establish whether BCR mutational load corresponds with cellular origin, we analyzed the mutational profiles of Bcl6 CD4 KO-derived and tdTomato^+^ cells in comparison to Bcl6 CD4 WT-derived cells. MBCs derived from Bcl6 CD4 WT mice, containing both GC-dependent and -independent cells, showed an average of 2.2 amino acid changes in the VH region. In contrast, Bcl6 CD4 KO GC-independent MBCs were almost fully unmutated, with an average of 0.3 amino acid changes in the VH region, while S1pr2-tdTomato^+^ GC-dependent MBCs exhibited a high mutational load, with an average of 7.1 amino acid changes in the VH region (**Figs. 3G, S4D**).

In sum, classical MBCs and a subset of atypical-like MBCs differentiate GC-dependently, while MZ-like and innate-like MBCs develop independently of the GC, consistent with their limited class-switching and reduced affinity maturation.

### GC-independent MBCs constitute a phenotypically distinct subset defined by a characteristic surface marker profile

Having confirmed the GC-independent origin of innate-like MBCs, we aimed to identify unique surface markers to enable their isolation and further study alongside classical, MZ-like, and atypical-like MBC populations. Since mRNA and surface protein expression are often poorly correlated (Li et al. 2020; Vallejo et al. 2022), we interrogated differential surface protein expression in the GC-dependent and-independent sc-sequencing dataset (**Figs. 3C, S4A–C**) and compared it to the expression of their corresponding genes. Using barcoded antibodies (Stoeckius et al. 2017), we found distinctive sets of surface proteins for the four MBC subsets, while only a few of their encoding genes showed correlated expression (**Fig. 4A**). Classical MBCs were characterized by high expression of the follicular marker proteins CD23 and CD55, while MZ-like MBCs expressed marginal zone markers CD21 and CD1d (**Fig. 4A, left**). This was in accord with the top cluster markers we identified by gene expression (**Fig. 1C**), which showed similar expression patterns (**Fig. 4A, right**). Conversely, innate-like and atypical-like MBCs were negative for both CD23 and CD21, but expressed several shared surface proteins (**Fig. 4A, left**), among them CD11b and CD29 (**Figure 4B**). While CD73 was identified as a strong marker for atypical-like MBCs through surface protein expression, its corresponding gene, *Nte5,* showed poor expression. Similarly, surface expression established CD11b as a robust marker of both innate-like and atypical-like MBCs, while *Itgam* expression was almost undetectable (**Fig. 4A, right**).

**Figure 4.**
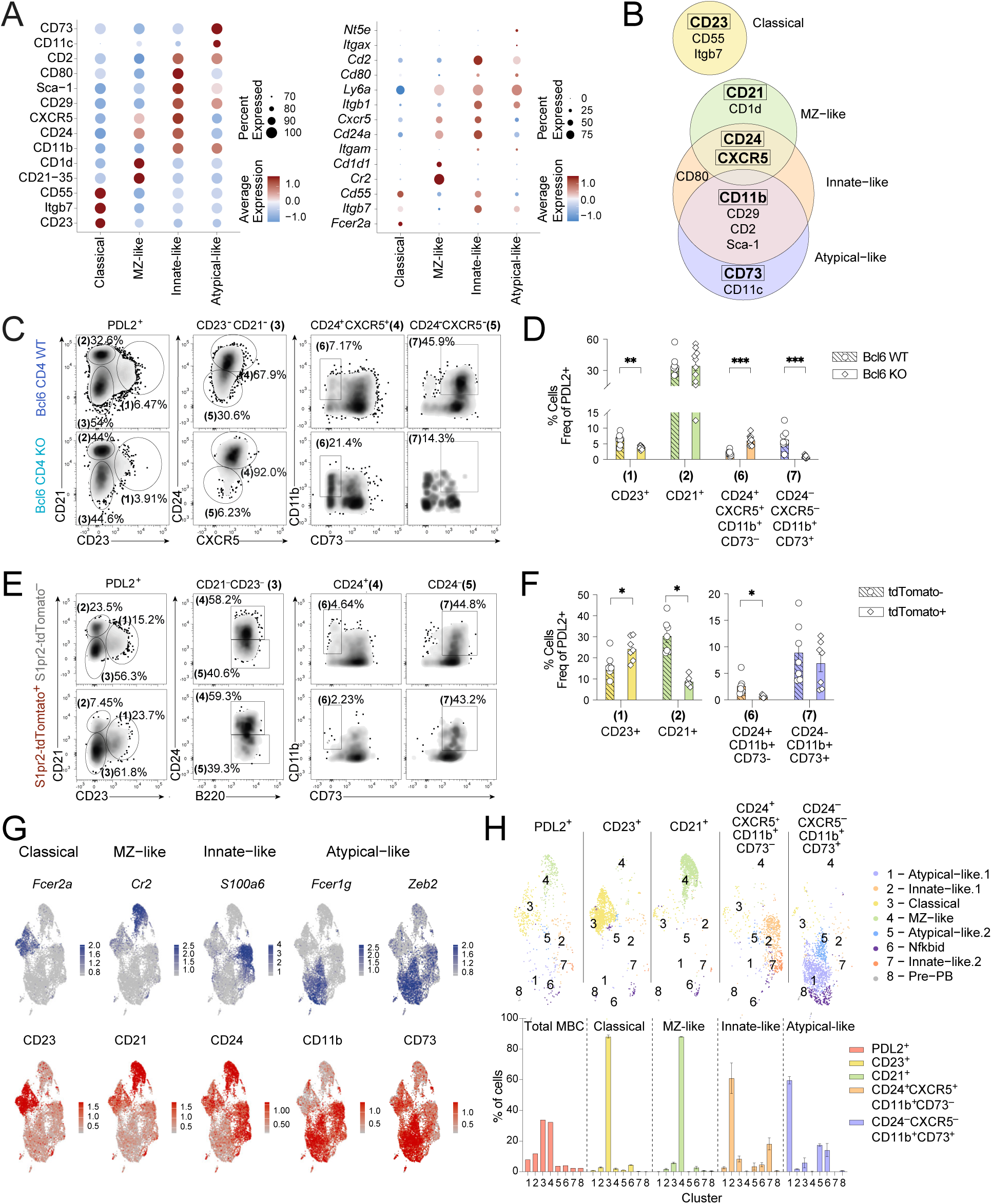
Identification of surface protein markers for isolation of MBC subsets. (A) Dot plot of differentially expressed surface markers (*left*) and their corresponding genes (*right*) obtained from the CITE-seq dataset (Fig. 3) comparing S1pr2-tdTomato^+^ and Bcl6 KO MBCs across classical, MZ-like, innate-like and atypical-like subsets. Dot size indicates the percentage of cells expressing each marker, and color intensity reflects scaled mean expression. (B) Euler diagram showing overlap of surface marker expression per subset. Framed surface markers were used to label and isolate the respective MBC populations by flow cytometry. (C) Density plots showing representative flow cytometry staining of MBCs (CD138^−^CD38^+^IgD^−^PDL2^+^). Candidate protein markers were fluorescently labelled in Bcl6 CD4 WT and Bcl6 CD4 KO mixed bone marrow chimeras. Mice were immunized i.p. (day 0) with 50 µg NP-CGG/AHG, boosted (day 28) with 50 µg NP-CGG/PBS and sacrificed 6–10 weeks after boost. Parent population indicated above each density plot. Percentages indicate frequency relative to the parent population. (D) Bar plots showing the quantification of MBC populations in Bcl6 CD4 WT (circles) and Bcl6 CD4 KO (diamonds) mixed bone marrow chimeras. Frequency of MBC populations defined by candidate surface protein markers identified in (A and B). CD23^+^: classical; CD21^+^: MZ-like; CD21^−^CD23^−^CD24^+^CXCR5^+^CD11b^+^CD73^−^: innate-like; CD21^−^CD23^−^CD24^−^CXCR5^−^CD11b^+^CD73^+^: atypical-like MBCs. Numbers in parenthesis indicate population hierarchy shown in density plots (C). Data is representative of two independent experiments. Statistical significance was determined using an unpaired Mann-Whitney test with Holm-Sidak correction for multiple comparisons. (E) Representative density plots displaying tdTomato^−^ and tdTomato^+^ MBCs from S1pr2 Cre^ERT2^ tdTomato mice immunized with 50 µg NP-CGG/AHG (day 0) and injected i.p. with 2 mg tamoxifen (days 4–7), boosted (day 28) with 50 µg NP-CGG/PBS and sacrificed 4–8 weeks after boost. Parent population indicated above each density plot. Percentages indicate frequency relative to the parent population. (F) Bar plots showing quantification of MBC populations in tdTomato^−^ (circles) and tdTomato^+^(squares). MBC populations defined by candidate surface protein markers identified in (A and B). CD23^+^: classical; CD21^+^: MZ-like; CD21^−^CD23^−^CD24^+^CD11b^+^CD73^−^: innate-like; CD21^−^CD23^−^CD24^−^CD11b^+^CD73^+^: Atypical-like MBCs. Numbers in parenthesis indicate population hierarchy shown in density plots. Data is representative of two independent experiments. Statistical significance was determined using a two-tailed Wilcoxon matched-pairs signed-rank test. (G) Feature plots displaying *top*: gene expression of *Fcer2a*, *Cr2*, *S100a6, Fcer1g and Zeb2*; and *bottom*: expression of surface markers CD23, CD21, CD24, CD11b, and CD73 on MBCs in UMAP. Scale: normalized UMI (Unique Molecular Identifier) counts. Cluster 9 was removed for visualization. (H) *Top*: UMAPs showing MBCs split by sorting strategy as indicated above each embedding. MBC populations were sorted using candidate surface proteins as indicated in (D). *Bottom*: Bar plots showing frequency of cells per cluster in each sorting strategy used. Sorting strategies were conducted with two biological replicates, except for PD-L2⁺ cells, which were assessed in a single replicate. In D and F, each symbol represents one mouse and mean ± SEM are marked. *p < 0.05, **p < 0.01, ***p < 0.001, and ****p < 0.0001.

Given that surface protein detection by barcoded antibodies is expected to correlate more closely with fluorescent antibody readouts, we selected combinations of markers to be tested by flow cytometry based on the differential surface protein analysis. We hypothesized that classical MBCs would be identifiable by CD23 expression, while we expected MZ-like MBCs to express high levels of CD21. Since innate- and atypical-like MBCs differed in their expression of CD73, CD24, and CXCR5 (**Fig. 4A–B**), we postulated that innate-like MBCs should be identifiable through expression of CD24, CXCR5, and CD11b, while atypical-like MBCs should be defined by CD73 and CD11b expression (**Fig. 4B**).

To test cluster-specific markers for the GC-derived classical and atypical-like MBC subsets by flow cytometry, we performed the NP-CGG prime-boost immunization in Bcl6 CD4 KO mice, which lack GC-derived MBCs, with Bcl6 CD4 WT mice as controls, as above (**Figs. 3A, S3A–B**). Total MBCs were identified by gating CD38^+^IgD^−^PDL2^+^ as previously described (**Fig. 1B**). Overall, total MBC frequency did not significantly differ between Bcl6 CD4 WT and Bcl6 CD4 KO mice (**Fig. S5A**). However, a CD23^+^ population (gate 1; proposed classical) was significantly decreased (by ∼42%) in GC-deficient Bcl6 CD4 KO mice relative to Bcl6 CD4 WT (**Fig. 4C–D**). Additionally, the CD24^−^CXCR5^−^CD11b^+^CD73^+^ population (gate 7; proposed atypical-like) showed a ∼97% reduction in GC-deficient Bcl6 CD4 KO mice compared to control mice (**Fig. 4C–D**). In contrast, the CD21^+^ population (gate 2; proposed MZ-like) did not significantly differ between WT and KO, while CD24^+^CD11b^+^CD73^−^ (gate 6; proposed innate-like) cells increased by ∼3-fold in GC-deficient Bcl6 CD4 KO mice. These findings, together with the differential gene and surface protein expression (**Figs. 4A–B, S4A**), suggest that the CD23^+^ and CD24^−^CXCR5^−^CD11b^+^CD73^+^ surface-marker combinations can identify classical and atypical-like MBCs, respectively, by flow cytometry.

To test cluster-specific markers for GC-independent MZ-like and innate-like MBCs, we returned to the S1pr2-CreERT2 tdTomato fate-mapping immunization model, where these populations would be underrepresented among the strictly GC-dependent tdTomato⁺ cells relative to tdTomato^−^ cells. Mice were immunized and boosted with NP-CGG, and tamoxifen was administered on days 4 to 8 to permanently label GC-derived MBCs. A CD21^+^ B cell population (gate 2; proposed MZ-like) was significantly reduced among GC-experienced S1pr2 tdTomato^+^ MBCs relative to tdTomato^−^ MBCs (mean 8.7% vs 30.4%). Similarly, the CD24^+^CD11b^+^CD73^−^ population (gate 6; proposed innate-like) was significantly decreased in GC-experienced tdTomato^+^ MBCs compared to tdTomato^−^MBCs (mean 0.68% vs 2.57%) (**Fig. 4E–F**). In contrast, the CD23^+^ population (gate 1; proposed classical) was significantly more frequent in tdTomato^+^ cells, while the CD24^−^CD11b^+^CD73^+^ population (gate 7; proposed atypical) showed equal representation across tdTomato^+^ and tdTomato^−^ cells (**Fig. 4E–F)**. Overall, these results demonstrate that CD21 and the CD24^+^CD11b^+^CD73^−^ surface protein combination are compelling candidates for selective labelling of MZ-like and innate-like MBCs, respectively.

To verify that our curated surface-marker combinations delineate the MBC clusters identified by gene-expression and enable the recovery of highly enriched subsets, we sorted the four MBC populations using our established markers and included a PDL2^+^CD38^+^IgD^−^ control sample encompassing all MBC subsets (**Fig. S5B**). Sc-sequencing on the sorted cells demonstrated that gene and surface markers identified for the respective MBC populations showed co-expression (**Fig. 4G**). Faceting the UMAP embeddings by sorting condition demonstrated high accordance of our gating strategy with transcriptional clustering (**Fig. 4H, top**), confirming that CD23 and CD21 identified the classical and MZ-like MBC subsets, respectively, while CD24 and CXCR5 labelled the two innate-like MBC clusters present in the dataset (**Fig. 4H, top**), with these populations being enriched to ∼80% purity (**Fig. 4H, bottom**). Furthermore, we also found that CD73 and CD11b surface expression resolved two atypical-like clusters (**Fig. S5C–E**); however, the gating scheme also captured *Nfkbid*^+^ MBCs, indicating heterogeneity within the sorted population (**Figs. 4H, S5C–D**), with enrichment of atypical-like populations to ∼60% purity (**Fig. 4H, bottom**). Overall, this isolated population did accumulate over time (**Fig. S5F**), which is a characteristic of atypical MBCs (Yi Hao et al. 2011; Du et al. 2019). Notably, alternative surface-marker combinations did not yield pure MBC subsets (**Fig. S5G**). Therefore, all subsequent analysis used the described gating schemes (**Figs. 4C–H**), prioritizing purity at the expense of excluding cells from specific clusters.

In summary, we identified surface-protein combinations uniquely expressed by each of the four MBC populations and validated their correspondence to the transcriptionally defined clusters to enable robust flow-cytometric identification and sorting of pure populations for further characterization.

### Innate-like MBCs arise independently of follicular and marginal zone extrafollicular B cell responses

Having established that innate-like MBCs arise independently of GC, we hypothesized an extrafollicular (EF) origin involving marginal zone (MZ), follicular B cells (FoB), or B1 cells. To test this, we used CD19Cre^+/+^ (CD19-KO) mice, in which CD19 deficiency abrogates MZ and B1 cell development while leaving FoB cell differentiation intact (**Figs. S6A–C**), restricting extrafollicular responses to the FoB compartment. WT and CD19-KO mice were NP-CGG prime-boosted, and MBC differentiation was evaluated by flow cytometry 60 days after primary immunization **(Fig. 5A**). Overall, CD19-KO mice showed a significant ∼5.7-fold increase in classical MBC frequency relative to WT controls, while MZ-like, innate-like and atypical-like MBCs were reduced by ∼5.0-, 3.8-, and 6.5-fold, respectively (**Figs. 5B–C**). This reduction in GC-independent MBCs was consistent with the near complete loss of MZ and B1 B cells that persisted in CD19-KO mice after immunization (**Fig. 5D**). Consequently, there was a ∼2.3-fold reduction in total MBCs, and we also confirmed the expected abrogation of the GC-response (∼90% reduction) (**Fig. 5E**). These results suggest that classical MBCs can differentiate from FoB during the EF response or from any remaining GC (**Fig. 5E**), while innate-like, MZ-like and atypical-like MBCs require precursors beyond the FoB compartment, implicating MZ or B1 cells as their progenitors.

**Figure 5.**
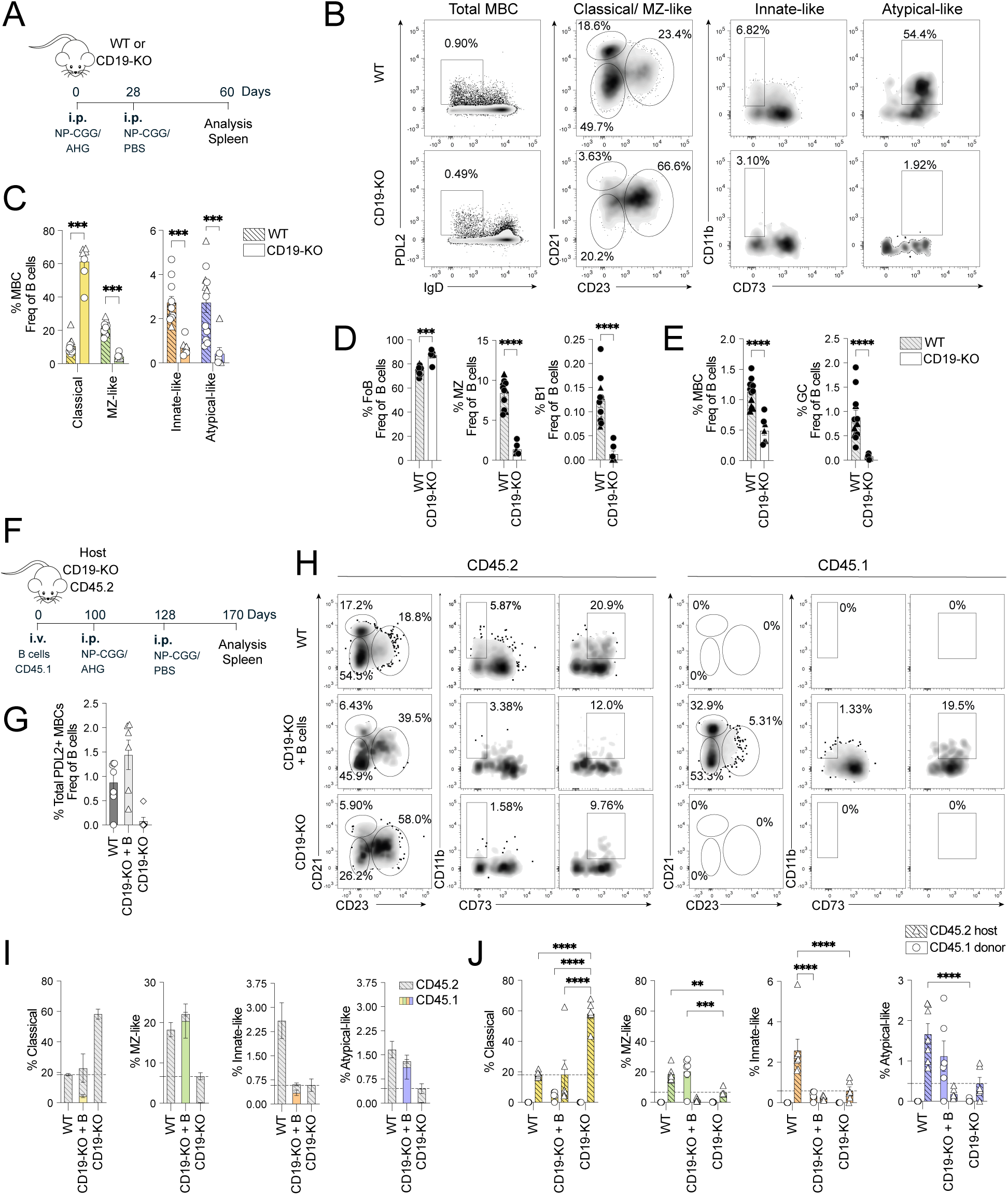
Innate-like MBCs do not develop from B2 follicular or MZ B cells. (A) Schematic of mouse immunization and boost model used. (B) Representative density plots of total MBCs and gated MBC populations in WT (*top*) and CD19-KO (*bottom*) mice. (C) Bar plots showing quantification of MBC subsets indicated on x-axes. Data shown was obtained from two independent experiments, mean ± SEM. (C, D and E): Circles indicate mice from experiment 1 and triangles mice from experiment 2. Significance was assessed using unpaired non-parametric Mann-Whitney tests or multiple t-test with Holm-Sidak correction. (B–E) Two experiments representative of three independent experiments are shown. (D) Bar plots showing B cells from WT (grey hatched bars) and CD19-KO (white solid bars) mice. Quantification of (left to right): splenic follicular B2 (FoB), MZ B cells, and B1 cells in peritoneal cavity after NP-CGG immunization. Due to lack of CD19 expression in CD19-KO animals, B1 B cells were gated as CD38^+^B220^lo^ (**Fig. S5B**). (E) Bar plots showing total MBCs (*left*) and GC frequency (*right*) of WT (grey hatched bars) and CD19-KO (white solid bars) mice after NP-CGG immunization. (F) Schematic of marginal zone reconstitution in CD19-KO mice, immunization and boost regimen. (G) Frequency of total PDL2^+^ MBCs in WT control (circles), CD19-KO transferred with CD45.1 B cells (triangles), and CD19-KO control (diamonds) mice. (H) Representative density plots of MBC populations from CD45.2 host (*left*) and CD45.1 B cell transferred (*right*). From top to bottom: WT control; CD19-KO transferred with CD45.1 B cells; CD19-KO control. (I) Stacked bar plot showing cumulative frequency of the indicated MBC populations. Hatched bars indicate MBC frequencies of CD45.2 recipients, solid bars indicate MBC frequencies of CD45.1 donor cells. (J) Bar plot showing CD45.1 and CD45.2 MBC frequencies in each experimental group. (I–J) Dashed horizontal lines show the average MBC frequency in negative control CD19-KO mice (MZ-like, innate-like and atypical-like MBCs) or positive control WT mice (classical MBCs) accordingly. Data is representative of three independent experiments. Statistical significance was determined using an unpaired Mann-Whitney test with Holm-Sidak correction for multiple comparisons. In all bar plots, each symbol represents one mouse, mean ± SEM is shown, *p < 0.05, **p < 0.01, ***p < 0.001, and ****p < 0.0001.

To determine whether MZ reconstitution was sufficient to restore MBCs in these three populations, we used a well-described CD19-KO MZ reconstitution system (Arnon et al. 2013; Liu et al. 2024), in which CD45.1 splenic B cells are transferred into congenic CD19-KO recipients (**Fig. 5F**). Once reconstitution of the MZ compartment was verified 8–16 weeks after transfer (**Fig. S6D**), we proceeded to NP-CGG prime-boost the mice (**Fig. 5F**). Upon immunization, total MBC frequency was restored to WT levels in MZ-reconstituted mice (**Fig. 5G**). MZ-like and atypical-like MBCs increased by ∼3.5- and 3-fold, respectively, relative to unreconstituted CD19-KO controls. Both populations were almost entirely composed of CD45.1 donor-derived cells and restored to WT-like frequencies, thus confirming a MZ origin (**Figs. 5H–I**). Classical MBC overrepresentation was corrected, decreasing by ∼2.5-fold compared to unreconstituted CD19-KO mice and returning to levels similar to WT controls (**Figs. 5H–I**). These findings suggest that MZ-niche reconstitution in CD19-KO mice largely restored B cell homeostasis across these populations, although atypical-like MBCs trended towards reconstitution without reaching significance (**Fig. 5J**). In contrast, MZ reconstitution did not restore innate-like MBCs, the frequencies of which remained significantly lower than in WT controls and indistinguishable from unreconstituted CD19-KO mice (**Figs. 5H–J**).

In conclusion, we confirmed that classical, MZ-like and atypical-like MBCs can originate from either FoB or MZ B cells during EF responses. In contrast, innate-like MBCs are unable to develop from these cell types alone.

### Innate-like CD24/CD11b-expressing MBCs are derived from B1 cells

As innate-like MBCs were not derived from FoB or MZ B cells during EF responses, B1 B cells remained as potential precursors of innate-like MBCs. We therefore investigated whether the transcriptomic identity of innate-like MBCs showed evidence of a B1 cell origin, and transcriptional signature analysis revealed that innate-like MBC clusters were enriched for B1 cell-associated genes (Luo et al. 2022) (**Figs. 6A–B).** B1 cells are also characterized by restricted V(D)J gene usage in their BCR repertoire. Therefore, we analyzed VH/VL gene usage across the four MBC populations and assessed their overlap with canonical B1 cell VH/VL pairs (Luo et al. 2022). We found that innate-like MBC displayed the most frequent usage of canonical murine B1 cell VH/VL pairs (mean = 31.5%) compared to classical, MZ-like, and atypical-like MBCs (mean = 0.2%, 0.3%, and 3.3%, respectively) **(Fig. 6C, Table S3**). Together, these results support a B1 origin for innate-like MBCs.

**Figure 6.**
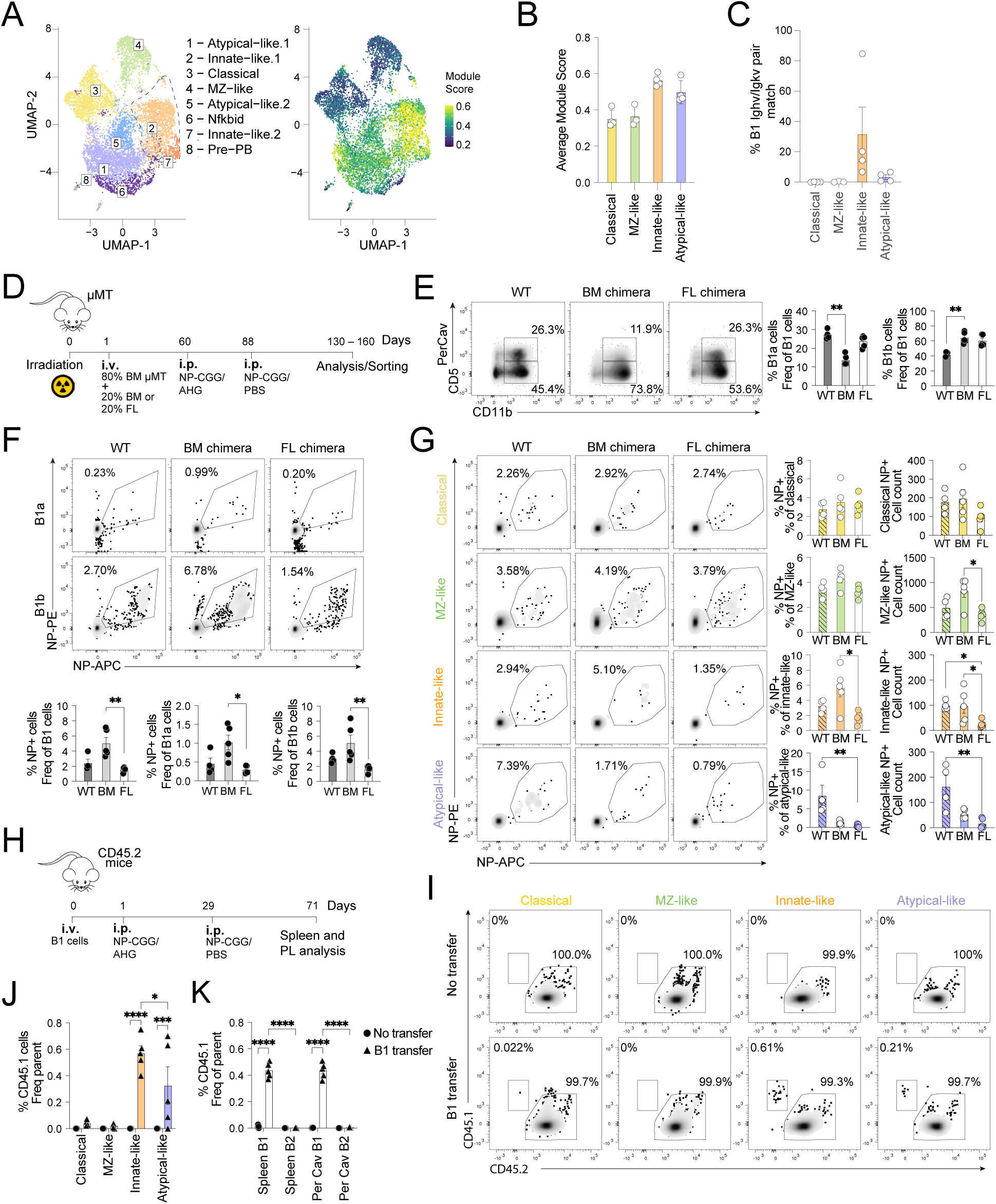
Innate-like MBCs derive from B1 B cells. (A) *Left*: UMAP from Fig. 4H, dashed outline indicates the innate-like MBC clusters. *Right*: Feature plot showing the B1 module score overlaid on the UMAP. The score calculated based on the genes listed in Table S2. Color scale indicates relative expression of the gene set in each cell. (B–C) MBC populations obtained from respective clusters of Seurat objects shown in Fig. 1 (clusters 1–4), Fig. 2 (clusters 1–4, 6) and Fig. 3 (clusters 1–5, 7) with each symbol representing one cluster. B1 VH/VL pairs used are detailed in Table S3. (B) Quantification of the average B1 module score per MBC population. (C) Frequency of B1 VH/VL pair match in each indicated subset. (D) Schematic of BM and FL chimera generation, and immunization protocol. (E) Representative density plots (*left*) and quantifications (*right*) of PerCav B1 populations gated as: 1) B1a: CD19^+^B220^lo^PDL2^-^CD11b^+^CD5^+^; 2) B1b: CD19^+^B220^lo^PDL2^-^CD11b^+^CD5^−^ in naïve WT, BM chimeric, or FL chimeric mice. (F) Representative density plots (*top*) and quantifications (*bottom*) of NP-specific B1 cells in PerCav of the indicated mice groups: WT, BM chimeras, or FL chimeras 6–10 weeks post NP-CGG boost. (G) Representative density plots (*left*), frequency (*middle*), and cell count (*right*) showing NP-specific cells in the indicated MBC subsets of NP-CGG immunized WT, BM chimeric, or FL chimeric mice. (H) Schematic of congenic B1 adoptive transfer model and immunization system. (I) Representative density plots highlighting the presence of CD45.1 adoptively transferred cells in the indicated MBC populations in two experimental groups of CD45.2 host mice: no transfer controls and B1-transferred mice. (J) Bar plots showing the frequency of CD45.1 cells in the indicated MBC subsets in B1 adoptively transferred or no-transfer control mice. (K) Bar plots showing the frequency of CD45.1 cells in the indicated spleen and PerCav B cell subsets in B1 adoptively transferred or no-transfer control mice. (E–G): Statistical significance was determined using a Kruskal–Wallis test followed by Dunn’s multiple comparisons test. (J–K): Statistical significance was assessed using a two-way ANOVA followed by a Tukey’s multiple comparisons test, with single pooled variance. In all bar plots, each symbol represents one mouse, mean ± SEM are shown, *p < 0.05, **p < 0.01, ***p < 0.001, and ****p < 0.0001.

To further assess whether innate-like MBCs arise from B1 cells, we leveraged the differential development of B1 cells: B1 cells primarily derive from fetal liver but are inefficiently generated from adult bone marrow, particularly B1a cells (Baumgarth 2017; Hayakawa et al. 2016). We therefore generated bone marrow (BM) and fetal liver (FL) chimeras by lethally irradiating µMT mice and reconstituting their hematopoietic compartment with either BM or FL cells from WT mice (**Fig. 6D**). Overall B1 cell frequency in the peritoneal cavity (PerCav) of chimeric mice was comparable to WT levels (**Fig. S7A**). Consistent with previous reports, BM chimeras displayed a ∼2-fold reduction in B1a cells and a ∼1.5-fold increase in B1b frequency relative to WT mice (**Fig. 6E**). Despite this baseline phenotype, total NP^+^ B1 cells were ∼3.4-fold more frequent in the PerCav of BM than FL chimeras under both naïve and NP-CGG immunized conditions. After immunization, PerCav B1a NP⁺ and B1b NP⁺ cells were ∼3.5- and ∼3-fold higher, respectively, in BM than in FL chimeras (**Figs. 6F, S7B–D**). A similar pattern was observed in the spleen, where NP-specific innate-like MBCs were increased by ∼3.5-fold in BM compared with FL chimeras in both frequency and absolute cell number. In contrast, the percentage of NP-specific classical and MZ-like MBCs was similar between BM and FL chimeras, while NP-specific atypical-like MBCs levels were reduced by ∼6.6 and ∼19.6-fold in BM and FL chimeras, respectively, compared with WT controls (**Fig. 6G**). Taken together, these findings reveal a correlation between NP-specific PerCav B1 cells and splenic NP-specific innate-like MBCs.

To directly establish the lineage relationship between innate-like MBCs and B1 cells, we employed a congenic cell tracking model. CD45.1^+^ B1 cells from the peritoneal cavity were adoptively transferred into CD45.2 recipient mice, which then underwent NP-CGG prime-boost immunization to induce MBC differentiation (**Fig. 6H**). At six weeks post boost, innate-like MBCs (mean = 0.57%) and to a lesser extent atypical-like MBCs (mean = 0.32%) were detected among CD45.1^+^ B cells in the B1 transfer recipients, representing the only MBC population with detectable B1-derived contribution (**Figs. 6I–J**). The frequency of donor-derived innate-like MBCs was consistent with the levels of donor-derived B1 cells detected in PerCav and spleen (**Fig. 6K**).

In time-course analysis, total MBCs increased by week 24 in both naive and immunized mice (**Fig. S7E**), indicating an immunization-independent expansion. In contrast, NP-specific MBCs were 10.2-fold higher 4 weeks after NP-CGG prime-boost and remained elevated throughout the time course (**Fig. S7F**). NP-specific innate-like MBCs and splenic B1 cells were also at higher frequencies in immunized mice relative to their unimmunized counterparts (**Fig. S7G**). To test whether this splenic accumulation reflected migration from the PerCav, we labeled B1 cells and NP^+^ innate-like MBC in the PerCav 6 weeks post-boost. B1 cells were significantly diminished in PerCav of immunized mice, while total innate-like MBCs remained unaltered (**Fig. S7H**). Concomitantly, splenic B1 cells and NP-specific innate-like MBCs—including class-switched NP^+^ cells—were ∼2-fold higher than naïve controls, while NP^+^ B1 cells did not change (**Fig. S7I**). Together, these reciprocal dynamics support a model of B1 cell migration from the PerCav to the spleen, followed by differentiation into NP-specific innate-like MBCs.

Collectively, these results demonstrate that the GC-independent innate-like CD24^+^CD11b^+^ MBC population originates from B1 cells, consistent with its B1-associated transcriptional signature and B1-biased VH/VL gene usage.

### Innate-like MBCs differentiate into antibody secreting cells and respond in an antigen-specific manner

Having established that innate-like MBCs derive from B1 B cells and are characterized by a limited VH/VL pairing, low affinity maturation, and low class switching, we sought to determine whether they nonetheless contribute to antigen-specific memory responses. To test whether innate-like MBCs differentiate into antibody-secreting plasma cells (ASCs) and mount NP-specific responses upon restimulation, we prime-boosted mice with NP-CGG and isolated the four splenic MBC populations (classical, MZ-like, innate-like and atypical-like), including CD5^+^ and CD5^−^ innate-like MBCs from PerCav, as well as B1a and B1b cells as their naïve PerCav counterparts. The isolated populations were stimulated *in vitro* on 40LB feeder cells which provide BAFF and CD40L, with addition of rmIL4 and CpG for four days (**Fig. 7A**) (Nojima et al. 2011). Flow cytometry analysis revealed that splenic innate-like, PerCav CD5^−^ innate-like MBCs and B1b cells proliferated the most, reaching approximately 6,000 cells per culture. Splenic innate-like and PerCav CD5^−^ innate-like MBCs generated the highest number of CD138⁺ plasmablasts, with ∼2,000 cells per culture (**Fig. 7B**). Collectively, these results indicate that innate-like MBCs showed enhanced plasmablast differentiation relative to the other MBC subsets.

**Figure 7.**
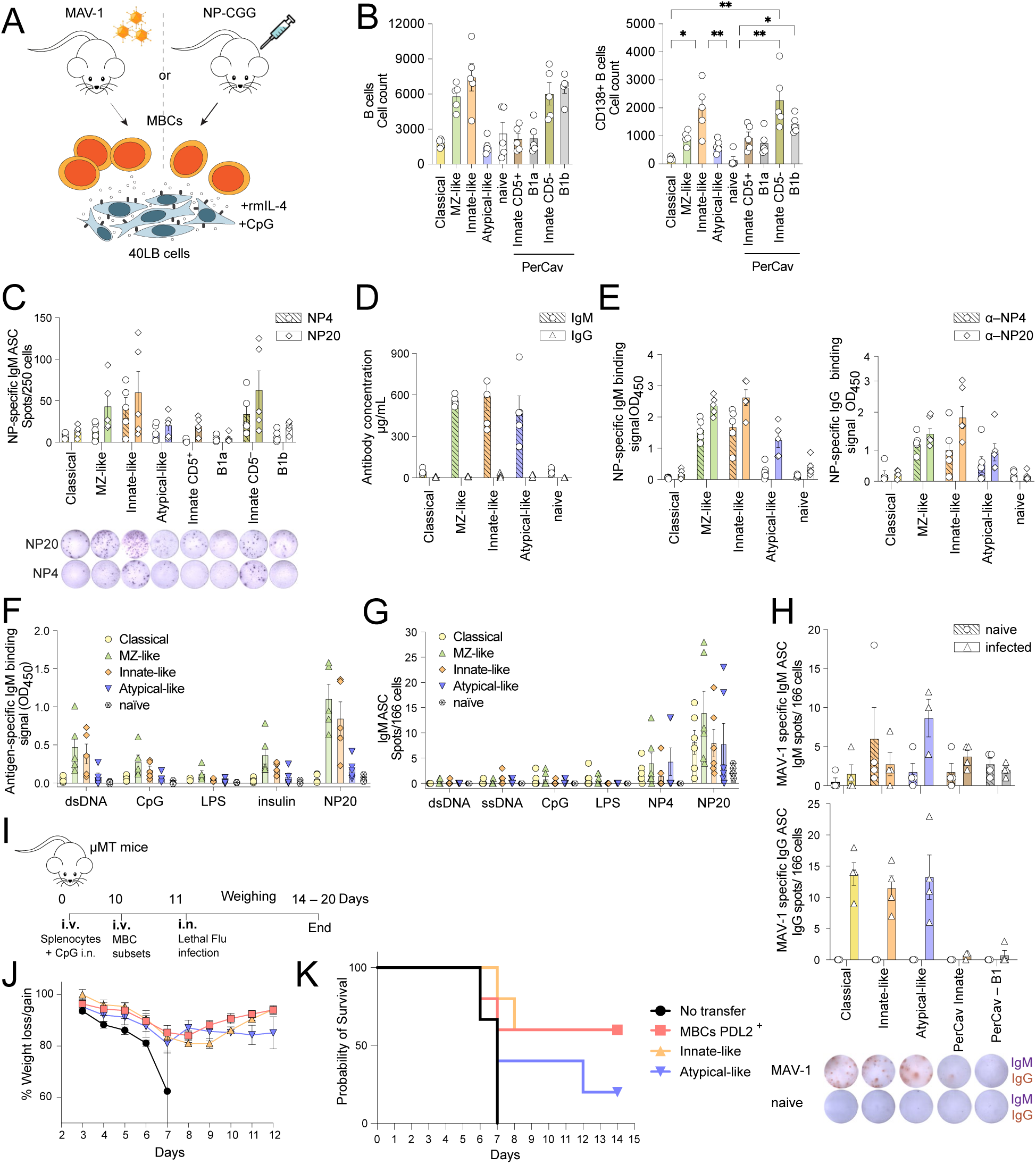
B1-derived innate-like MBCs differentiate into antigen-specific ASCs and confer protection from recurring infection. (A) Schematic of MBC culture using MBCs from MAV-1 infected or NP-CGG immunized mice. MBC populations were cultured for 4 days with 40LB feeder cells (Nojima culture) supplemented with rmIL-4 and CpG. (B–G). MBCs were obtained from NP-CGG primed/boosted mice. Indicated MBC populations were isolated as follows (*left* to *right*): 1) Classical, 2) MZ-like, 3) splenic innate-like and 4) atypical-like MBCs were sorted as described in Fig. 3; 5) naïve B cells: B220^+^CD19^+^CD38^+^IgD^+^; 6)PerCav Innate-like CD5^+^: CD19^+^B220^low^PDL2^+^CD11b^+^CD5^+^; 7) PerCav B1a: CD19^+^B220^lo^PDL2^-^CD11b^+^CD5^+^; 8)PerCav Innate-like CD5^−^: CD19^+^B220^low^PDL2^+^CD11b^+^CD5^−^; 9)PerCav B1b: CD19^+^B220^lo^PDL2^-^CD11b^+^CD5^−^. (B) Quantification of total cell counts by flow cytometry on Nojima culture day 4. *Left*: total B cells (B220^+^CD19^+^). *Right*: CD138^+^ plasmablasts. (C) Quantification of ELISPOT assays. MBCs were cultured as in (A), cells were recovered and seeded in antigen-coated plates for ELISPOT assay. *Top:* NP20-(diamond) and NP4 (circle)-specific spot counts of IgM are plotted. Spots counted per 250 cells seeded on day 0 of Nojima culture. Each symbol represents ASCs differentiated from one MBC population of an independent mouse. *Bottom:* Representative well images of each condition. Data from one representative out of 7 independent experiments. (D) Total IgM (circle, hatched bars) and IgG (diamond, solid bars) concentration (µg/mL) in the supernatant from the indicated MBC populations harvested on d4 of the Nojima culture. Data from one representative out of 3 independent experiments. (E) Antigen-specific IgM production assessed by ELISA using NP-coated plates. Binding signal reported as background-subtracted OD_450_. NP20-(diamond, solid bars) and NP4 (circles, hatched bars)-specific IgM (*left*) and IgG (*right*) binding signal in supernatants from the indicated MBC populations after four days of culture. Each symbol corresponds to an independent mouse. Data representative of 3 independent experiments. (F) Antigen-specific IgM detected by ELISA on day 4 supernatants using antigen-coated plates as indicated. Binding signal reported as background-subtracted OD_450_ for each MBC population (as indicated in the legend). Each symbol corresponds to an independent mouse. Data from one representative out of 2 independent experiments. (G) ELISPOT assay of IgM ASCs specific for the listed antigens. Plot shows spot counts per 166 cells seeded in Nojima culture. Each symbol corresponds to an independent mouse. (H) Quantification of ELISPOT assays. MBCs were obtained from mice infected with 1.2 x 10^3^ TCID_50_ MAV-1 i.p. and cultured as in (A). Cells were recovered at day 4 and seeded in antigen-coated plates for ELISPOT assay, MBC subsets were selected as (*left* to *right*): 1) Classical, 2) MZ-like, 3) splenic innate-like and 4) atypical-like MBCs were sorted as described in Figure 3; 5) PerCav Innate-like: CD19^+^B220^low^PDL2^+^; 6) PerCav B1: CD19^+^B220^low^PDL2^−^. Plots show ASCs spots count MAV-1 specific IgM (*top*) and IgG (*bottom*) per 166 cells seeded on d0 of the Nojima culture. Representative wells (*bottom*) show IgM (purple) and IgG (red) spots. Each symbol corresponds to an independent mouse. Data representative of 3 independent experiments. (I) Schematic of influenza protection assay. Briefly, bronchoalveolar lymphoid structures were induced in µMT mice at day 0 by transferring 1x10^5^ splenocytes i.v. and 15 µg CpG i.n. per mouse. After 10 days MBCs and CD4^+^ T cells were adoptively transferred into µMT mice and next day they received a lethal dose of influenza A (PR8) i.n. (900 PFU per mouse). (J) Body weight of mice following influenza infection over a 12-day period. Data are shown as percent of initial body weight for the indicated groups. Mice received 32,800 total PDL2^+^ MBCs, 8700 innate-like MBCs or 1900 atypical-like MBCs obtained from lung and spleen of previously infected mice. (K) Kaplan–Meier survival curves showing viability of mice following influenza infection over 12 days for the indicated groups.

Innate-like MBCs contained the highest number of NP^+^ cells, averaging ∼90 cells, whereas all other populations remained below 50 cells (**Fig. S8A**). To further evaluate NP-specific responses of the differentiated plasmablasts, we performed ELISPOT assays. Splenic innate-like and PerCav CD5^−^ innate-like MBCs produced the highest number of NP20-specific IgM ASCs (mean ∼60 per 250 cells), whereas MZ-like MBCs generated a lower response (mean of ∼40 per 250 cells). Nonetheless, all three populations exhibited higher numbers of NP-specific ASCs than classical, atypical MBCs and B1 cell controls (mean 5–20) (**Fig. 7C**). We also assessed high affinity responses using NP4 coating, which due to its low hapten density selectively captures high-affinity ASCs. Splenic innate-like and PerCav CD5^−^ innate-like MBCs produced the highest number of NP4-specific IgM ASCs (30–40) compared with the other B cell populations (2–14), indicating greater high-affinity antibody output (**Fig. 7C**). When ASC responses were normalized to plasmablasts numbers to account for differences in proliferation, all MBC populations exhibited increased ASC responses compared to B1a and B1b cells (**Fig. S8B**). However, classical MBCs generated the highest number of NP-specific IgM (∼18) and IgG (∼5) ASCs per 250 CD138^+^ cells, consistent with their enrichment for NP-binding cells (**Fig. 2C**). Altogether, this indicates that the high number of NP-specific ASCs found in the innate-like MBC culture is due to their robust proliferation and plasmablast differentiation (**Fig. 7B**).

Overall, class switched responses of differentiated ASCs were limited in this system (**Fig. S8C**). Accordingly, IgM concentration exceeded IgG in culture supernatants (**Fig. 7D**). Consistent with the high NP-specific ASCs obtained, supernatant from stimulated innate-like MBCs exhibited high NP20-specific IgM signal relative to the other populations.

Surprisingly, NP4-specific IgM signal was only slightly lower than NP20-specific IgM (**Fig. 7E**), indicating substantial high-affinity antibody production. A similar signal trend was observed for IgG accumulated in the supernatant despite the low concentration of total IgG detected (**Fig. 7D–E**). When antibodies in the supernatant were normalized to total IgM, NP-specific IgM binding signal was similar across all populations, while innate-like and atypical-like MBCs showed slightly higher NP-specific IgG binding signal (**Fig. S8D**). Taken together, these results demonstrate that innate-like MBCs efficiently differentiate into ASCs and are capable of mounting robust NP-specific responses upon restimulation.

B1 B cells are known to secrete polyreactive antibodies (Ehrenstein and Notley 2010; Mattos et al. 2024). To evaluate whether B1-derived innate-like MBCs also show polyreactivity, we performed ELISA binding assays and measured secreted antibodies against common antigens targeted by polyreactive antibodies (Tiller et al. 2007). Although innate-like and MZ-like MBCs exhibited modest signals of polyreactive IgM, NP-specific antibodies predominated (**Fig. 7F**). This was further confirmed by ELISPOT analysis, in which NP20-specific IgM ASCs averaged ∼15 per culture, whereas only ∼1 ASC per unrelated antigen tested showed polyreactivity (**Fig. 7G**).

Overall, innate-like MBCs undergo robust ASC differentiation upon restimulation, retain NP-specificity, and show limited polyreactivity, collectively demonstrating their functional effector capacity.

### Innate-like MBCs produce antiviral ASCs and provide protection against lethal influenza infection

After investigating the ASC responses of innate-like MBCs to model hapten antigen, we sought to evaluate their potential to contribute to protective immune responses in a physiologically relevant viral infection context. We first examined virus-specific ASC differentiation of MBCs that had been generated in response to mouse adenovirus 1 (MAV-1). On day 28 post-infection, splenic MBCs, PerCav innate-like MBCs and PerCav B1 cells were isolated and cultured as described above (**Fig. 7A**). ELISPOT analysis of cultured cells revealed that in contrast to hapten immunization, MAV-1 infection triggered a mostly class-switched ASC response, with only atypical-like MBCs showing an increased number of virus-specific IgM ASC (∼8 per culture) in cells derived from infected mice (**Fig. 7H**). All splenic MBC populations produced a robust MAV-1 specific IgG response, averaging ∼13 IgG ASCs per culture, while PerCav innate-like MBCs remained mostly unresponsive (**Fig. 7H**). Since the spleen is a peripheral site of MAV-1 replication (Kajon, Brown, and Spindler 1998; Moore, Brown, and Spindler 2003), the preferential accumulation of MAV-1 specific MBCs in the spleen may indicate a local or resident nature. Thus, splenic innate-like MBCs differentiated into MAV-1-specific ASCs, produced class-switched antibodies, and accumulated at a site of viral proliferation.

Having confirmed that innate-like MBCs are able to robustly differentiate into anti-viral ASCs, we next aimed to test their capacity to protect against infection using a previously established influenza assay (Onodera et al. 2012). We adoptively transferred naïve splenocytes from WT mice into µMT recipients and administered CpG intranasally to induce bronchus-associated lymphoid structures. After 10 days, MBCs isolated from lungs and spleens of mice previously infected with influenza A PR8 were adoptively transferred into µMT recipients. Recipient mice were then infected with a lethal dose of PR8, weight was monitored daily, and mice were euthanized upon reaching 20% of initial mass loss (**Fig. 7I**). We observed that weight loss peaked between 8–9 days post-infection, after which surviving mice started to recover (**Fig. 7J**). Strikingly, while atypical-like MBC produced only 20% protection, mice which received innate-like MBCs showed 60% survival, comparable to positive control mice which received total PDL2^+^ MBCs (**Fig. 7K**). Thus, innate-like MBCs retain virus-specific memory and confer robust protection against lethal viral rechallenge.

## DISCUSSION

In this study, we implemented a multifaceted approach to analyze MBC populations and identified a previously uncharacterized B1-derived innate-like MBC population that, while detected in previous gene expression studies (Gregoire et al. 2022; Riedel et al. 2020; Risley et al. 2025), has not been experimentally explored. Our results set the B1-derived innate-like MBCs apart as an individual branch of the overall memory response, increasing MBC diversity by contributing the B1 BCR repertoire, which differs from the B2 repertoire (Baumgarth 2011). Our findings are consistent with prior observations that the MBC compartment shows higher clonal diversity than PCs, and that GC B cells have a reduced clonal overlap with MBCs generated in parallel (Purtha et al. 2011; Shinnakasu et al. 2016; Viant et al. 2020; Wong et al. 2020).

B1 cells are widely identified as an early barrier of protection, producing what is often referred to as “natural antibodies”, which have antigen-specificity against common pathogenic structures or self-antigens encoded in their germline V genes (M. Boes 2000; Casali and Schettino 1996; Palma et al. 2018). This enables B1 B cells to provide rapid protection during initial pathogen encounter, before adaptively mediated immune responses are fully engaged (Marianne Boes et al. 1998; Baumgarth 2016; Baumgarth et al. 2000). While the role of B1 B cells in memory responses had not been well elucidated, studies have shown that they confer protection to the T-independent pathogens *Francisella tularensis (Cole et al. 2009)*, *Borrelia hermsii* (Alugupalli et al. 2004), and *Salmonella typhi* (Marshall et al. 2012). In these studies, B1 cells were able to mediate fast-acting but also long-lasting protection, including from recurring infection, reminiscent of memory responses. These earlier studies correlated the presence of antigen-specific B1 cells and protective serum IgM levels. Similarly, earlier research showed sustained plasma cells and antibody production by antigen-specific B1b cells in response to T-independent type 2 immunogen (Hsu et al. 2006), hinting to a memory like response. Our findings demonstrate that B1 cells effectively give rise to innate-like MBCs in response to T-dependent antigen immunization and infection. Furthermore, this response involved both IgM and IgG production and included detectable levels of higher-affinity antibodies, particularly following hapten immunization, and under these conditions, innate-like MBCs produced more antigen-specific ASCs than B1 cells. These findings distinguish innate-like MBCs from B1 cells and provide an alternative perspective to previous research in which long-term cross-pathogen protection was primarily attributed to B1 cell responses to pathogens inducing T-independent immunity (Alugupalli et al. 2004). Similarly, previous work showed that mice lacking a PDL2^+^ B1a subpopulation were more susceptible to sepsis, identifying PDL2 expression on B1 cells as protective, though the identity of this population beyond its surface phenotype remained unresolved (Guo et al. 2019). Our findings indicate that PDL2 is a distinguishing marker of CD24^+^CD11b^+^innate-like MBCs, suggesting that the protective population identified by Gou et al. may in fact represent a memory B cell subset not previously recognized as such. Expanding on this premise, our data suggest that innate-like MBCs may contribute to MBC clonal diversity, potentially enhancing the breadth of recall responses against pathogen variants. In addition, consistent with the canonical role ascribed to B1 cells, innate-like MBCs may function as an early “natural” barrier during recurrent infection by rapidly differentiating into antibody-secreting cells upon antigen encounter. Since their repertoire is enriched for specificities against previously encountered pathogens compared with naïve B1 cells, the antibodies they secrete contain a higher fraction of antigen-specific antibodies. These antibodies could be present both locally at sites of infection or viral replication and systemically, given that B1 cells are major contributors to serum antibody levels (Choi et al. 2012).

Previous studies of B1 long-term protection observed that memory formation required either B1a or B1b cells, depending on the pathogen or antigen (Alugupalli et al. 2004; Cole et al. 2009; Haas et al. 2005; Hsu et al. 2006; Marshall et al. 2012). This aligns with evidence that B1a and B1b cells differ in their repertoires, with B1b cells exhibiting a broader repertoire (Cunningham et al. 2014; Srikakulapu et al. 2022; Tornberg and Holmberg 1995) and the capacity to accumulate somatic mutations (Roy et al. 2009). This notion is supported by our findings that B1b cells are the main responders in the T-dependent NP-CGG immunization model. However, it remains to be elucidated whether innate-like MBCs are preferentially generated from B1a or B1b cells.

B1 cells are predominantly located in the pleural and peritoneal cavities, and they transiently relocate to secondary lymphoid tissues during infection in mice and humans (Shen et al. 2025; Waffarn et al. 2015). Similarly, in this study, we characterized innate-like MBCs in the peritoneal cavity, lungs, and spleen. The presence of antigen-specific innate-like MBCs in both PerCav and spleen following hapten immunizations suggests that these could either migrate between tissues or arise locally from peritoneal or splenic B1 cells, respectively. Curiously, innate-like MBCs that differentiated after MAV-1 infection were exclusively present in splenic tissue, a site of MAV-1 replication (Kajon, Brown, and Spindler 1998), while no MAV-1-specific innate-like MBCs were found in the peritoneal cavity despite intraperitoneal administration of the virus. Similarly, influenza-specific innate-like MBCs were mainly found in the lungs, the main site of H1N1 replication. In line with this, Gregoire et al. identified three distinct MBC clusters in the lung after influenza infection, whose gene expression profiles were reminiscent of atypical-like, classical, and innate-like MBCs (Gregoire et al. 2022). These findings raise the possibility that innate-like MBCs establish tissue-residency determined by the type of immunization or pathogen replication.

In addition to B1-derived innate-like MBCs, we also consistently identified classical, MZ-like, and atypical-like MBCs by multiple methods. Interestingly, while the atypical-like MBCs are reminiscent of age-associated B cells and showed gene expression similarities to CD11c^+^Tbet^+^ MBCs, *Tbx21* expression in this subset was negligible. MBC isolation from GC fate-mapped S1pr2-tomato^+^ cells yielded a bona fide, fully GC-derived, and highly mutated *Tbx21*^hi^ CD11c^hi^ population that clustered separately from atypical-like MBCs. Atypical-like MBCs, as defined by our surface markers, constituted a heterogeneous population composed of multiple gene expression clusters and arising from three distinct ontogenetic origins: MZ, B1, and GC B cells. While Tbet^+^ MBCs have been extensively studied in recent years, there are some conflicting results regarding their origin (Knox, Myles, and Cancro 2019; Song et al. 2022). Therefore, a multi-cluster and multi-origin nature for atypical-like MBCs could explain some of these observed discrepancies; however, further analysis would be needed to elucidate differences within these subsets.

In summary, our findings reveal an expanded landscape of memory B cell types, including B1-derived innate-like MBCs, which are capable of somatic mutation and high-affinity recall responses. These cells represent a distinct arm of immunological memory that contributes to BCR diversification and may enhance the breadth of humoral immunity. Defining its role in vaccination and local immune protection will be an important area for future study.

### Limitations of the study

In this study, we identified MBCs as PDL2^+^IgD^-^, which allowed us to consistently isolate GC-independent MBC populations, as well as to largely recapitulate MBC types described in the literature. As discussed, while we did consistently identify an atypical-like MBC subset, we only isolated the bona fide Tbet^+^ MBCs when sorting fate-mapped GC-derived MBCs. As these cells expressed lower levels of PDL2 than the remaining MBC subsets, they may be excluded by our sorting scheme. A second subset of cells described in the literature but not recovered in significant numbers in this study were ISG^+^ MBCs (Cooper et al. 2024). We detected this subset in some experiments, indicating that it is captured by our selection scheme; however, because it is expanded during chronic viral infection, it is likely not efficiently generated by the hapten immunization used in our single-cell sequencing experiments.

While we used a combination of genetic models and congenic transfer experiments to demonstrate the B1-origin of the innate-like MBCs, we were not able to identify a mouse model with a specific B1 deficiency to demonstrate selective depletion of only innate-like MBCs. Finally, it remains for future investigations to fully elucidate whether innate-like MBCs are generated exclusively locally, or whether they migrate between tissues following immunization or infection.

## SUPPLEMENTAL TABLE LEGENDS

**Table S1.** Gene signature used for Tbet^+^CD11c^+^ MBCs module score analysis. Genes were derived from (Song et al. 2022) filtering genes adj p<0.001 and positive 2-fold log2FC. Module scores were calculated using the Seurat AddModuleScore function with default parameters.

**Table S2.** Gene signature used for B1 module score analysis. Genes were derived from (Luo et al. 2022) filtering genes adj p<0.001 and positive 2-fold log2FC. Module scores were calculated using the Seurat AddModuleScore function with default parameters

**Table S3.** List with B1 VH/VL pairs used to calculate frequency of B1 VH/VL and MBCs VH/VL pair match.

## Acknowledgements

We would like to thank all members of the Batista lab for their assistance with our experiments, as well as the Ragon Flow Cytometry Core. We thank Deepak Rao for providing bone marrow from Bcl6 CD4 WT and Bcl6 CD KO mice. We also thank Anastasia Yandulskaya-Blue of the Ragon Scientific Editing Platform for improvements to this text. ChatGPT was used as an editing tool to develop the Abstract and Claude in the Materials and Methods to expedite obtaining IDs for the reagents list.

## Declarations of Interest

FDB has consultancy relationships with Adimab, Third Rock Ventures, and *The EMBO Journal*, and founded BliNK Therapeutics.

## Funding

This research was funded by the Gates Foundation (INV-046626 to F.D.B.), flexible funds from The Ragon Institute of Mass General Brigham, MIT, and Harvard (to F.D.B.), and European Molecular Biology Organization (EMBO) Postdoctoral Fellowship EMBO ALTF 482-2019 (to M.B.).

## MATERIAL AND METHODS

### Experimental animals

All mouse lines were obtained from The Jackson Laboratory (Bar Harbor, ME). Bones from Bcl6^fl/fl^CD4 Cre^−/–^ and Bcl6^fl/fl^CD4 Cre^+/–^ were obtained from Deepak Rao, Brigham and Women’s Hospital. Mouse breeding, housing, maintenance, and experimental procedures were performed at the animal facility of either the Mass General Brigham Center for Comparative Medicine (CCM) or the Vivarium of the Ragon Institute of Mass General Brigham, MIT, and Harvard. For experiments, male and female mice between 8–36 weeks old were used. Up to four mice per cage were housed under standard 12h light/dark cycle, with room temperature at 22°C, and free access to food and water. All animal experiments were conducted in accordance with the Institutional Animal Care and Use Committee (IACUC) of Massachusetts General Hospital (MGH)’s approved Animal Study Protocols 2016N000286 and 2016N000022. The MGH CCM and Ragon Vivarium are Association for Assessment and Accreditation of Laboratory Animal Care (AAALAC) International-approved programs.

### NP-CGG immunization

Animals were immunized via intraperitoneal (i.p.) injection of 50 µg NP_43_-CGG (Biosearch Technology) in Alhydrogel (InvivoGen). To study primary responses, animals were culled at day 7 and spleens harvested to analyze B cells using flow cytometry. To induce memory responses, mice were injected i.p. with 50 µg NP_43_-CGG (Biosearch Technology) in Alhydrogel (InvivoGen) and 4 weeks later boosted i.p. with 50 μg NP-CGG in PBS. Spleen and peritoneal lavage were harvested 60–100 days after secondary immunization and B cell responses were analyzed by flow cytometry, ELISPOT, and ELISA.

### Infection

Influenza A (H1N1 A/PR/8/34) was obtained from ATCC (VR-95). For Influenza A infection mice were anesthetized with isoflurane. Anaesthetized mice were then intranasally inoculated with 50–900 PFU of Influenza virus A/Puerto Rico/8/1934 (PR8) H1N1 strain in 20 µl PBS. To induce memory B cell response doses in the lower range (50–500 PFU) were used. MBCs were analyzed by flow cytometry or purified by cell sorting 2 to 4 months after inoculation. High doses (900 PFU) of influenza were used for lethal infection of mice during protection assays. For the protection experiments, uMT mice received 5x10^6^ C57BL/6 splenocytes intravenously (i.v.) and 15 µg of CpG ODN-1826 (Invivogen) intranasally (i.n.) in PBS (Gregoire et al. 2022; Onodera et al. 2012). After 10 days, mice received isolated MBCs sorted from lungs of previously infected C57BL/6 mice (low dose 2 to 4 months prior). The day after, mice were challenged with 900 PFU of Influenza A PR8 virus, weight was measured daily, and mice were euthanized if they exhibited ≥ 20% loss of initial mass.

Mouse Adenovirus-1 (MAV-1) was obtained from ViraQuest (North Liberty, IA, USA). For MAV-1 infections, mice were intraperitoneally infected with 1.2x10^3^ TCID50/mouse. MBCs were harvested 30 days post-infection.

### Cell harvest from tissues and tissue preparation

Animals were culled and lymphoid organs collected. To generate cell suspensions from soft organs, organs were passed through a 70 μm cell strainer in PBS supplemented with 2% FBS (FACS buffer). Lung suspensions were obtained with the mouse lung dissociation kit (Miltenyi) according to manufacturer instructions. The suspended cells were counted on a LUNA automated Cell Counter (Logos Biosystems, Anyang, KR) and kept on ice until further manipulation.

For flow cytometry analysis of fresh organs, red blood cells in spleen, peritoneal lavage, or lungs were lysed with 1X Erythrocyte lysis buffer (ACK Gibco or Lonza) 3 minutes at 4°C and washed with FACS buffer.

To harvest the peritoneal lavage, 5 mL PBS was injected into the intact peritoneal cavity and gently massaged. Then, lavage was extracted using a sterile Pasteur pipette and transferred into a 48-well deep well plate at 4°C until further manipulation.

For bone marrow extraction, femurs and tibias were removed and cleaned. Bones ends were cut with scissors, and bone marrow was extracted by centrifugation into FBS containing Eppendorf. For flow cytometry analysis bone marrow was extracted by smashing one bone in FACS buffer using a mortar and pestle. After both types of bone marrow extraction, cells were filtered through 70 μm cell strainer.

Fetal liver was obtained from day 21 embryos after decapitation. Fetal livers were pooled together and smashed using a 70 µm cell strainer. Cells were counted and stored in FBS supplemented with 10% DMSO at −80°C until further use.

Blood was collected from tail veins in heparin-coated tubes. Red blood cells were lysed twice using red blood cell lysis buffer (Gibco) for 5 minutes at 4°C. After lysis, white blood cells were kept at 4°C. For serum collection cheek bleeds were performed and blood collected in serum separator tubes (SST).

### Flow cytometry and cell sorting

Single-cell suspensions from blood, spleen, lymph nodes, peritoneal lavage, or bone marrow were prepared. Fc receptors were blocked with anti-CD16/32 antibody (eBiosciences) diluted to 5 µg/mL in PBS and, simultaneously, dead cells were stained using the LIVE/DEAD Fixable Dead Cell Stain Kit (Thermo Fisher Scientific, Waltham, MA, USA) for 15 minutes at 4°C. After washing, cells were stained with fluorescently labelled antibody cocktails for 20 minutes at 4°C. When using biotinylated antibodies, cell suspensions were washed following antibody labeling and incubated for 10 minutes on ice with labeled streptavidin. Before acquisition cells were filtered through a Multiscreen-Mesh Filter Plate, 60 µM ((MilliporeSigma, Burlington, MA, USA). For cell counts 25 µl of CountBright Absolute Counting Beads (Thermo Fisher Scientific, Waltham, MA, USA) were added per well. Data were acquired on a BD flow cytometer Symphony A5 Cell Analyzer (Becton, Dickinson and Company, Franklin Lakes, NJ, USA).

For intracellular staining, cells were first surface stained as described before, and then resuspended in fixation/permeabilization buffer (BD Cytofix/Cytoperm™ kit, BD Bioscience) and incubated 30 minutes at 4°C. After fixation, cells were washed with PermWash (BD Bioscience) and antibodies diluted in PermWash were added and incubated 30 minutes at 4°C. Next, cells were washed twice with PermWash. After the last wash, cells were resuspended in FACS buffer and analyzed by flow cytometry.

Cells were sorted using a 70µM nozzle. For scRNA-seq, cells were sorted into a PCR tube containing 5–20 µL PBS supplemented with 10% FBS. Cells were then loaded onto the 10x Genomics Chromium Controller targeting up to 40,000 cells per sample. For MBC or B1 transfer experiments cells were sorted using a 70 µM nozzle into Eppendorf tubes containing PBS supplemented with 10% heat inactivated mouse serum were collected in house.

### NP labelling

Activated APC or PE (Agilent) and NP-e-Aminocaproyl-Osu (Biosearch Technologies) were mixed in PBS for 2 hours at room temperature in a total volume of 40 μL. Fluorophore and NP were prepared at different ratios ranging from 40:1 to 2:1. The different mixtures were then diluted to a final volume of 200 μL and dialyzed against PBS at 4°C. To test conjugation efficiency, splenocytes from NP-CGG immunized animals and non-immunized controls were stained and analyzed by flow cytometry.

### MZ reconstitution and adoptive transfers

For marginal zone B cell reconstitution, spleens were collected from donor mice with a C57BL/6J (CD45.1^+/+^) background. Spleens were crushed through a 70 μm cell strainer and subjected to pan B cell isolation kits (Miltenyi Biotec). Isolated B cells were then quantified for live cells on a LUNA-FX7 automated cell counter (Logos Biosystems) and adjusted to the desired number and volume (200 μl/mouse) in phosphate-buffered saline (PBS) before transfer into CD45.2^+/+^ recipient mice by i.v. injection through tail vein. Approximately 1.2 x 10^7^ B cells were injected (Arnon et al. 2013) except where otherwise specified. Between 8–12 weeks mice were culled and MZ reconstitution assessed by flow cytometry.

For B1 cell adoptive transfers, peritoneal lavage was collected from CD45.1^+/+^ mice and pooled together. Total B1 cells were sorted as B220^low^CD19^+^ and 1x10^5^–2 x10^5^ B1 cells i.v. injected through the tail vein.

### Bone marrow and fetal liver chimeras

Bone marrow or fetal liver recipient animals were given acidified water at all times. To generate chimeras, μMT or RAG1-KO host mice were lethally irradiated with 5 Gy twice with a gap of 4 hours between these 2 doses. To reconstitute the depleted BM niche, 24 hours later mice were i.v. injected with a total of 2 x 10^6^ cells. For wildtype BM or fetal liver chimeras, a mixture of 75–80% μMT bone marrow cells and either 20–25% bone marrow or fetal liver cells were injected into irradiated uMT recipients. For Bcl6^fl/fl^ CD4 Cre chimeras, irradiated RAG1-KO hosts were i.v. injected with a mixture of 75% RAG1-KO BM and either 25% Bcl6^fl/fl^ CD4 Cre^−/–^ (WT) or Bcl6^fl/fl^ CD4 Cre^+/–^ (KO) BM. All chimeras were evaluated eight weeks later for bone marrow niche reconstitution by analyzing CD4+ T cells and B220^+^ B cells in blood samples by flow cytometry.

### *In vitro* Nojima culture

Fibroblasts expressing CD40L and BAFF, referred to from now on as 40LB cells (Nojima et al. 2011), were cultured and maintained in Dulbecco’s modified Eagle’s medium (DMEM) supplemented with 10% fetal bovine serum (FBS), 10mM HEPES (pH 7.3), 1mM sodium pyruvate, 100 U/ml penicillin/streptomycin, 400 µg/mL geneticin G418 and 5 µg/mL puromycin (feeder cell medium). To be used in B cell co-cultures, 40LB cell were grown until 80% confluency and treated with Mitomycin C (10 µg/mL, Sigma) for 2 hours at 37°C, 5% CO_2_ to stop cell proliferation. After incubation cells were washed 3 times with PBS at room temperature and detached from flasks by trypsinization. Treated 40LB cells were then counted and 1x10^5^ cells per well were seeded into 96-well plates. Cells were incubated overnight at 37°C, 5% CO_2_ in feeder medium to allow cell attachment. Next day, cells were placed in 100 µL complete B cell medium supplemented with CpG (6 µg/mL) and IL4 (2 ng/mL). MBC were added on top the 40LB layer in 100 µL fresh B cell medium and co-cultured for 4 days at 37°C, 5% CO_2_. On day 4, B cells were collected into deep well plates, spun down, supernatant was collected, and cells were resuspended in one volume of B cell media for further use.

### ELISA

Polystyrene high bind microplates (Corning) were coated with NP20-BSA or NP4-BSA diluted in PBS (1 µg/mL, Biosearch Technologies) at 4°C overnight for NP-CGG immunizations. The next day, plates were washed 3 times in PBS 0.01% Tween 20 (Sigma) and blocked for 2 hours at room temperature with FACS (PBS + 2% FBS) buffer. After blocking, supernatant from Nojima cultures was added and incubated at 4°C overnight. The plates were then washed 3 times in PBS-0.01% Tween 20 and incubated with IgM-HRP or IgG-HRP (1 µg/mL in FACS buffer, SouthernBiotech) for 2 hours at room temperature. Then, plates were washed 3 times in PBS-0.01% Tween 20. TMB substrate (Thermo Fisher Scientific) was added, and color development was monitored before stopping the reaction with 1 M H₂SO₄. The absorbance was measured at 450 nm with a SpectraMax iD3 plate reader (Molecular Devices).

To measure total IgM and IgG in supernatant Mouse IgM Uncoated ELISA Kit and Mouse IgG (Total) Uncoated ELISA Kit (Thermo Fisher Scientific) were used following manufacture’s protocols.

### ELISPOT

Enzyme-linked immunosorbent spot (ELISPOT) multiscreen plates (Millipore) were activated with 99% ethanol (Sigma) for 1 minute. Plates were washed 3 times with PBS and coated overnight with NP_20_-BSA at 4°C. Plates were then washed with PBS 3 times in sterile conditions and blocked for at least 2 hours at 37°C with complete B cell medium. Single-cell suspension of spleens or bone marrow were then plated at different cell densities and incubated for 24 hours at 37°C in complete B cell medium. Plates were then washed 3 times using PBS-T and incubated with HRP conjugated anti-IgM or IgG (Southern Biotech) for 1 hour. Finally, plates were washed with PBS-T and developed with TMB solution (Sigma). Reactions were stopped by rinsing the wells with water. The dried plates were stored at room temperature and the number of spots quantified using an ELISPOT counter (Cellular Technology Limited). For dual color ELISPOT once 24 hour incubation was finished, plates were washed 3 times using PBS-T and incubated with HRP conjugated anti-IgM and AP conjugated anti-IgG (Southern Biotech) for 1 hour. After incubation, plates were washed with PBS-T and developed first with BCIP/NIST (Sigma; prepared according to manufacturer’s instructions) for 5 min then washed with tap water 3 times, tap dried and incubated with TMB solution for 5 min. This reaction was stopped with water. The dried plates were stored at room temperature and the number of spots quantified using an ELISPOT counter (Cellular Technology Limited).

### 10x Single-cell sequencing

Single-cell RNA-seq libraries were prepared using the 10x Genomics Chromium Next GEM Single Cell 3′ kit v3.1 or 5′ kit v2 according to the manufacturer’s instructions (10x Genomics, Pleasanton, CA, US). For datasets generated using the 5′ chemistry, paired B cell receptor (BCR) libraries were prepared in parallel. Cell hashing with hashtag oligonucleotides (HTOs) was performed in all experiments, and antibody-derived tags (ADTs) were included in selected experiments. Libraries were quantified using the Tapestation 4200 (Agilent, Santa Clara, CA) and the Qubit fluorometry assay (AAT Bioquest, Sunnyvale, CA), and sequenced on the NextSeq2000 sequencer (Illumina, San Diego, CA).

Raw sequencing data were processed using Cell Ranger (10x Genomics). Gene expression libraries were processed using Cell Ranger v6.0.0, v7.0.0, or v7.2.0, and V(D)J libraries using Cell Ranger v6.1.0, v7.0.0, or v7.2.0. For datasets processed with Cell Ranger v7.0.0 and later, gene expression and BCR libraries were jointly processed using the Cell Ranger multi pipeline. All datasets were aligned to the mm10-2020-A reference (10x Genomics). Downstream analyses were performed in Seurat v4.3.0 (Yuhan Hao et al. 2021). Using established QC parameters (Tirosh et al. 2016), we set the following exclusion criteria: cells with fewer than 200 detected genes, >8% mitochondrial gene expression, or fewer than 50 detected housekeeping genes were excluded. HTO-based demultiplexing was performed using Seurat’s HTODemux function.

Gene expression data were normalized using either LogNormalize or SCTransform. Immunoglobulin variable (V) genes were excluded or regressed out prior to running FindVariableFeatures. Dimensionality reduction was performed using principal component analysis (PCA) and cells were visualized using Uniform Manifold Approximation and Projection (UMAP).

For datasets comprising multiple experiments, integration was performed using Harmony to correct for batch effects while preserving biological variation. Graph-based clustering was conducted using the Louvain algorithm, with resolution parameters specified in the corresponding figure legends.

For experiments incorporating ADTs, protein counts were normalized using centered log-ratio (CLR) transformation and analyzed in parallel with transcriptomic data.

BCR datasets were aligned to the corresponding 10x Genomics reference genome for mouse (vdj_GRCm38_alts_ensembl-5.0.0 or vdj_GRCm38_alts_ensembl-7.0.0).

Module scores were calculated using AddModuleScore function in Seurat with default parameters, gene lists used to calculate score are available in Tables S1 and S2.

### Statistical analysis

Sample sizes were chosen based on prior work in which similar phenotypic characterization and similar defects had been reported; no data was excluded. When data was normally distributed groups were compared with parametric Student’s t-test and P-values were calculated using PRISM software. When distribution was not normal or N too small, data was compared with nonparametric Student’s t-test with Welch’s correction. For multiple measurements performed for two experimental groups, data was compared using two-way ANOVA. Statistically significant differences are denoted on the figures as: \**P*≤0.05, \*\**P*≤0.01, \*\*\**P*≤0.001, \*\*\*\**P*≤0.0001.

## KEY RESOURCES TABLE

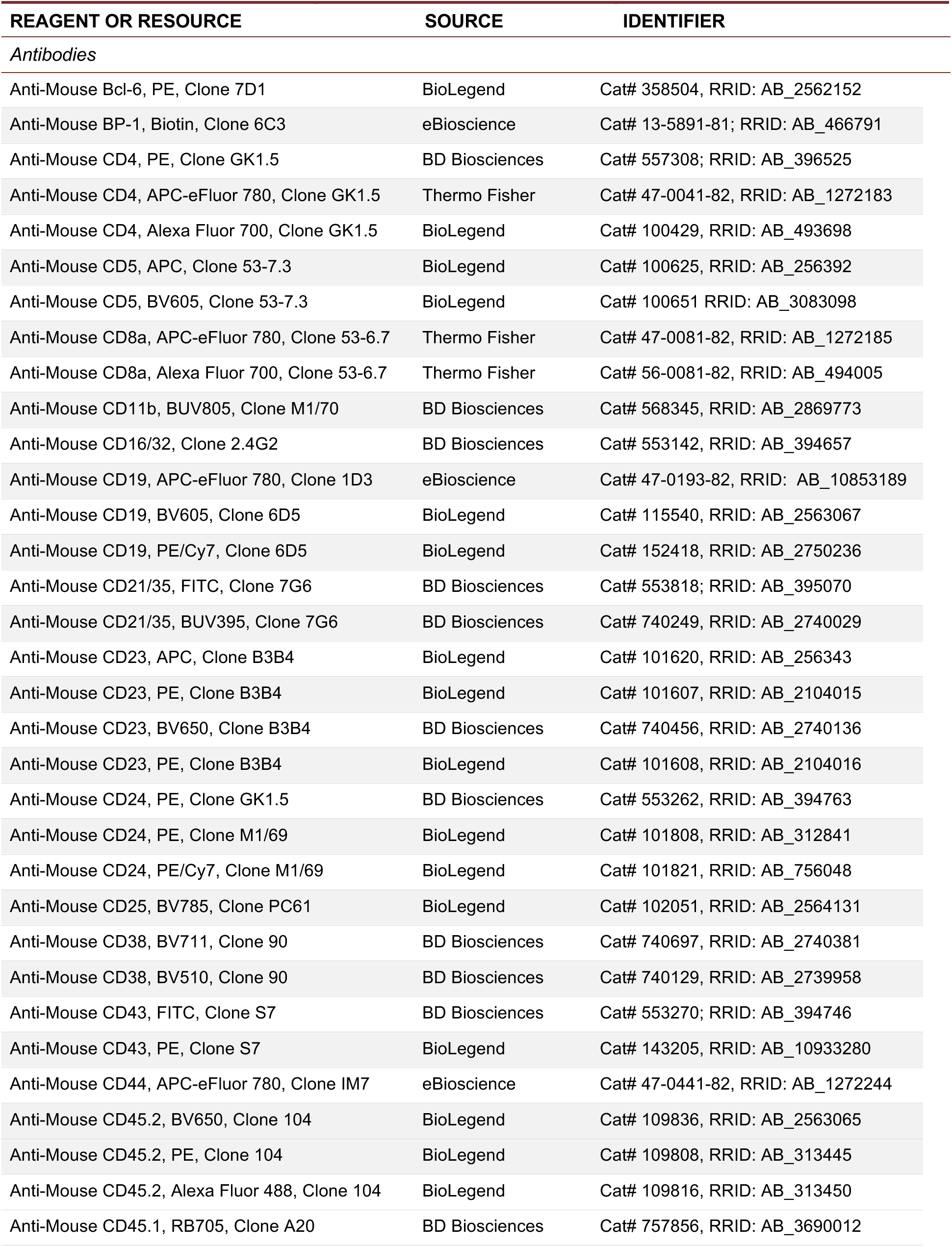
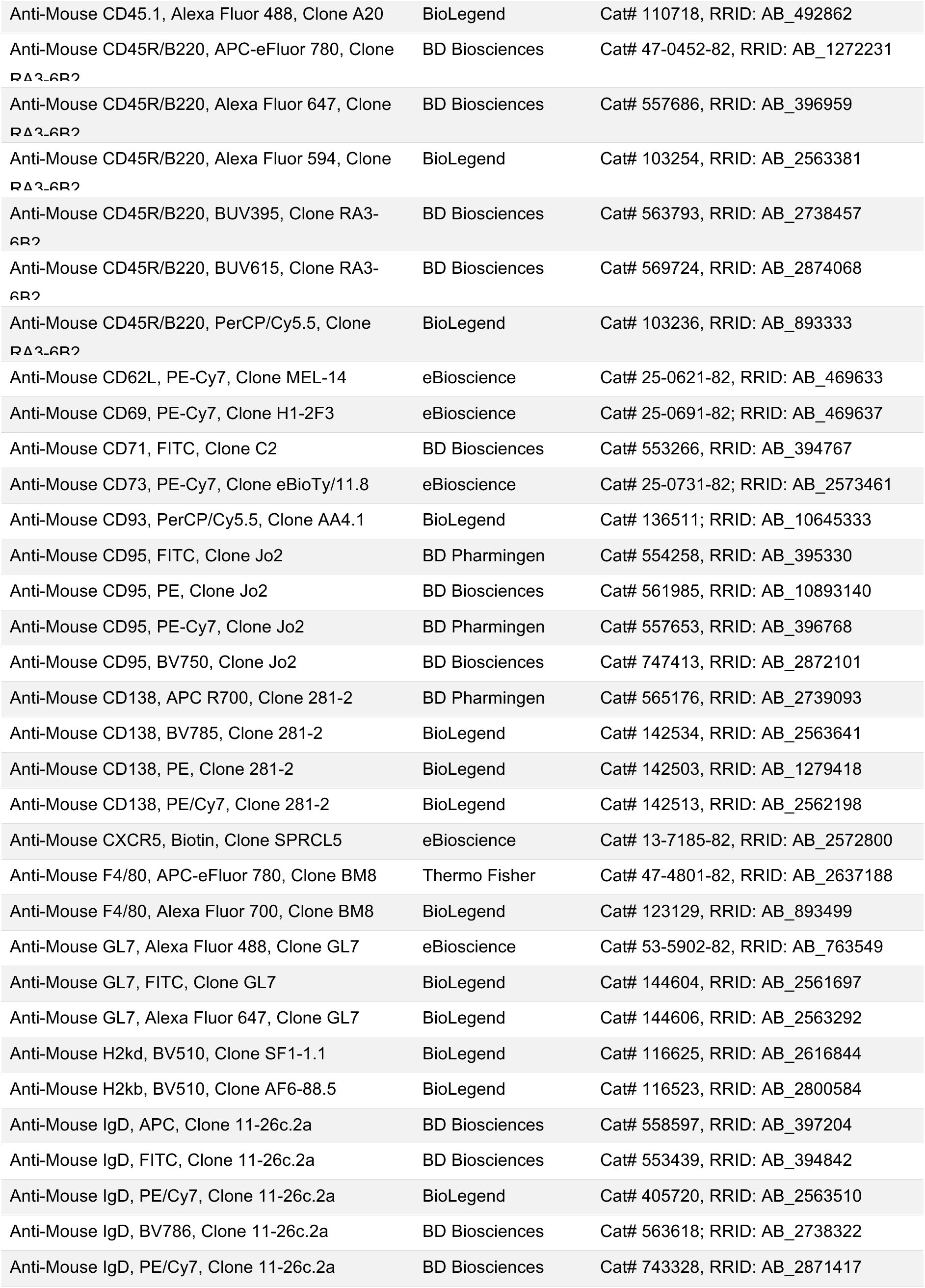
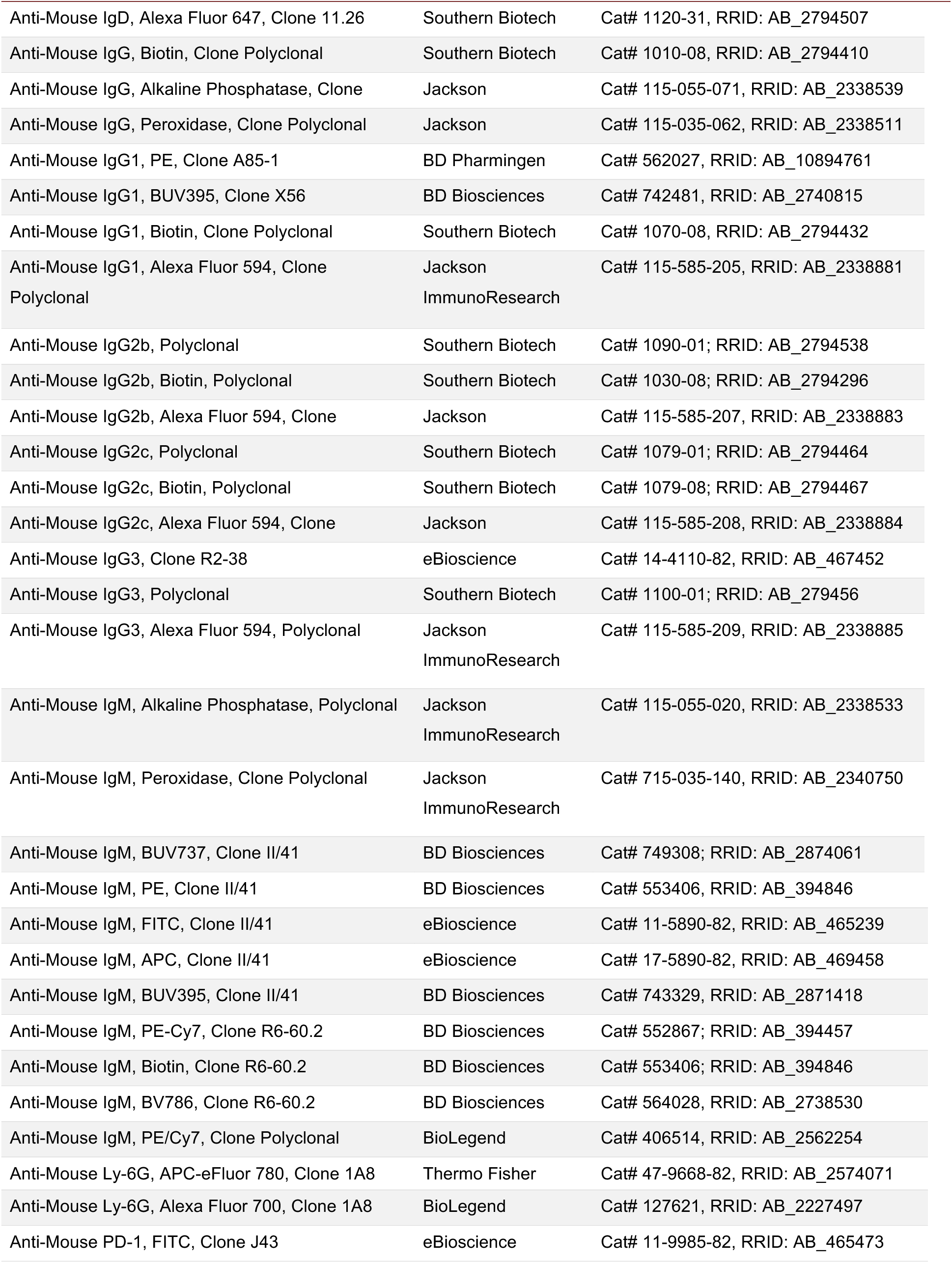
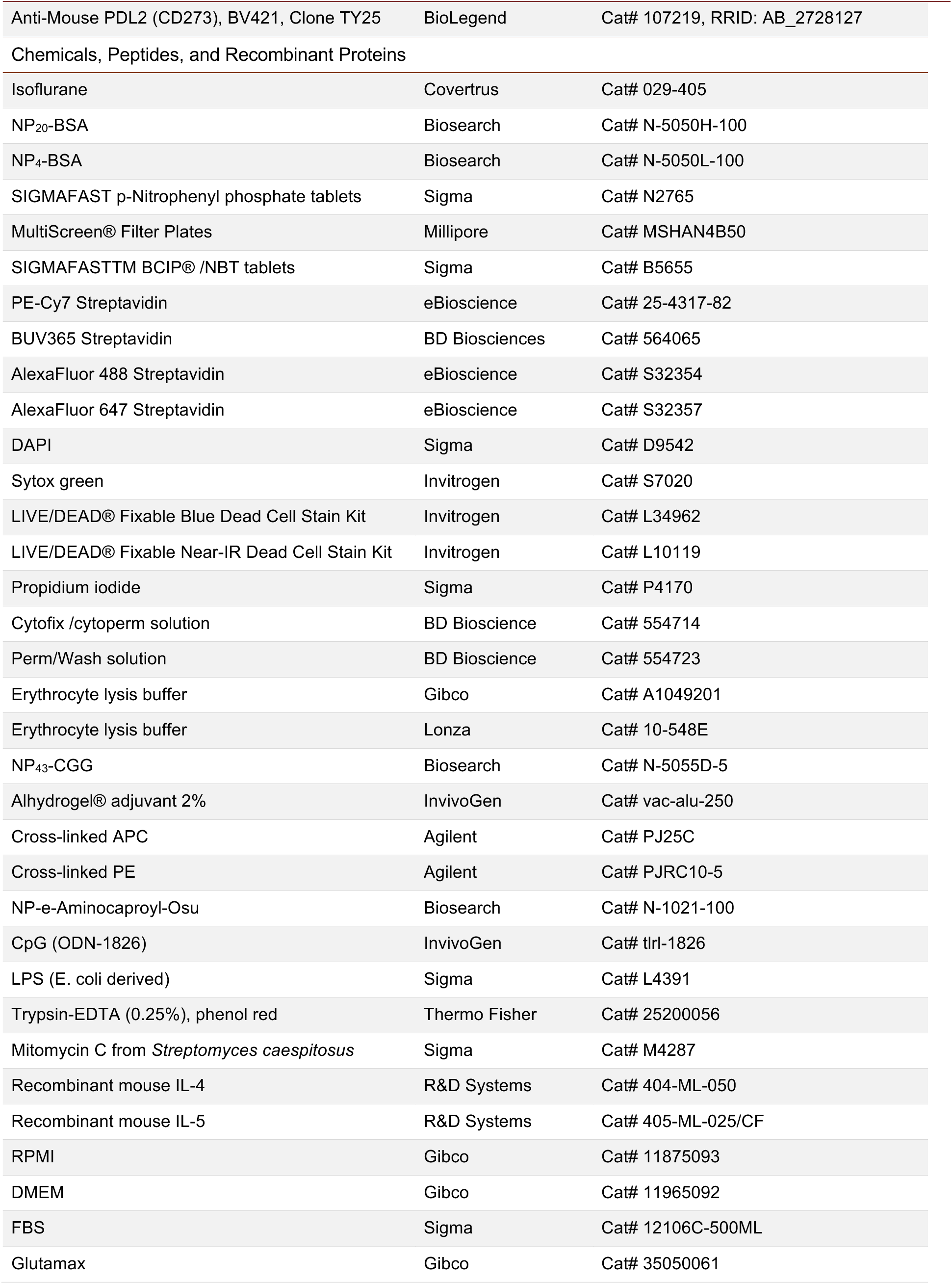
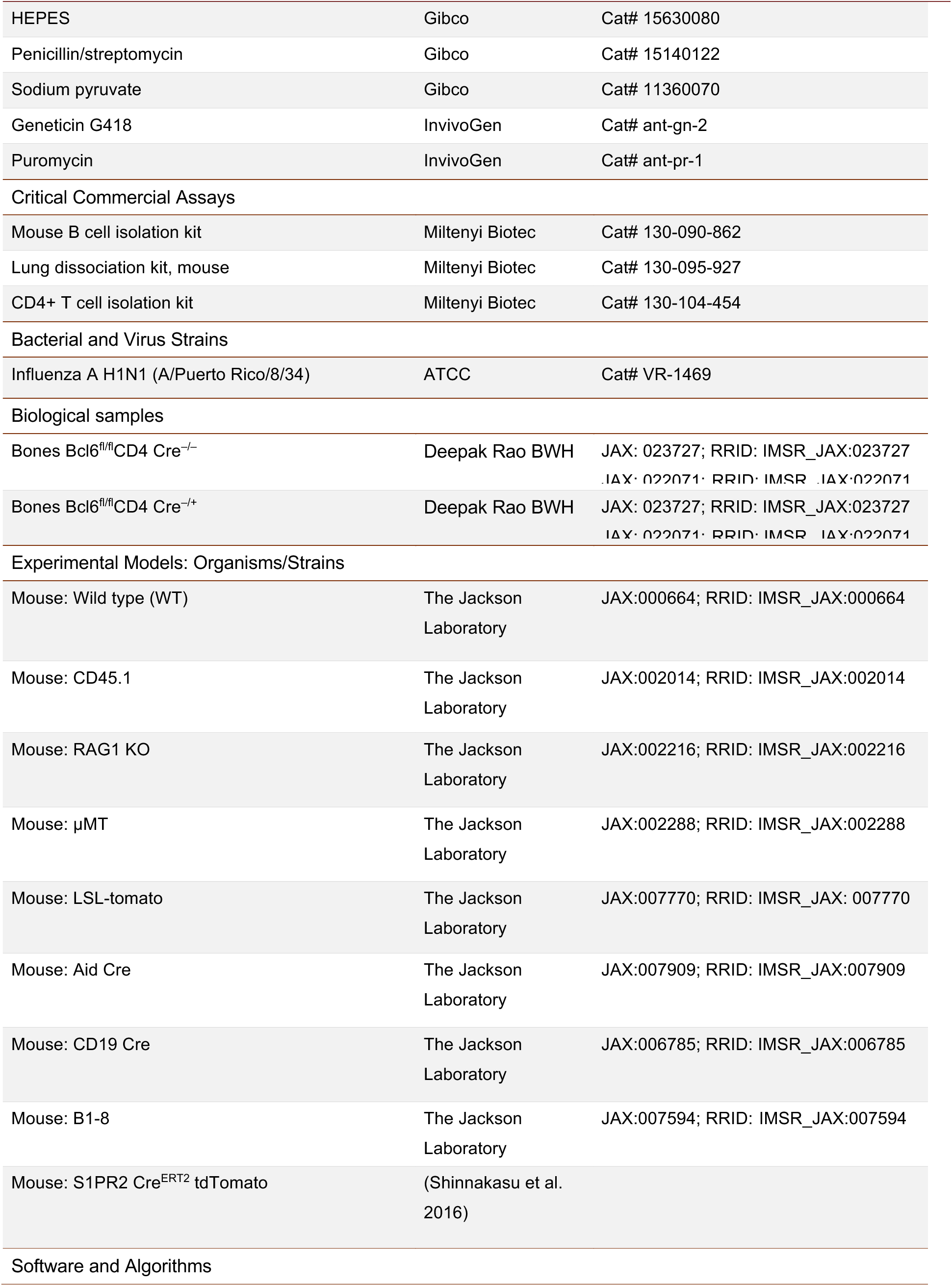
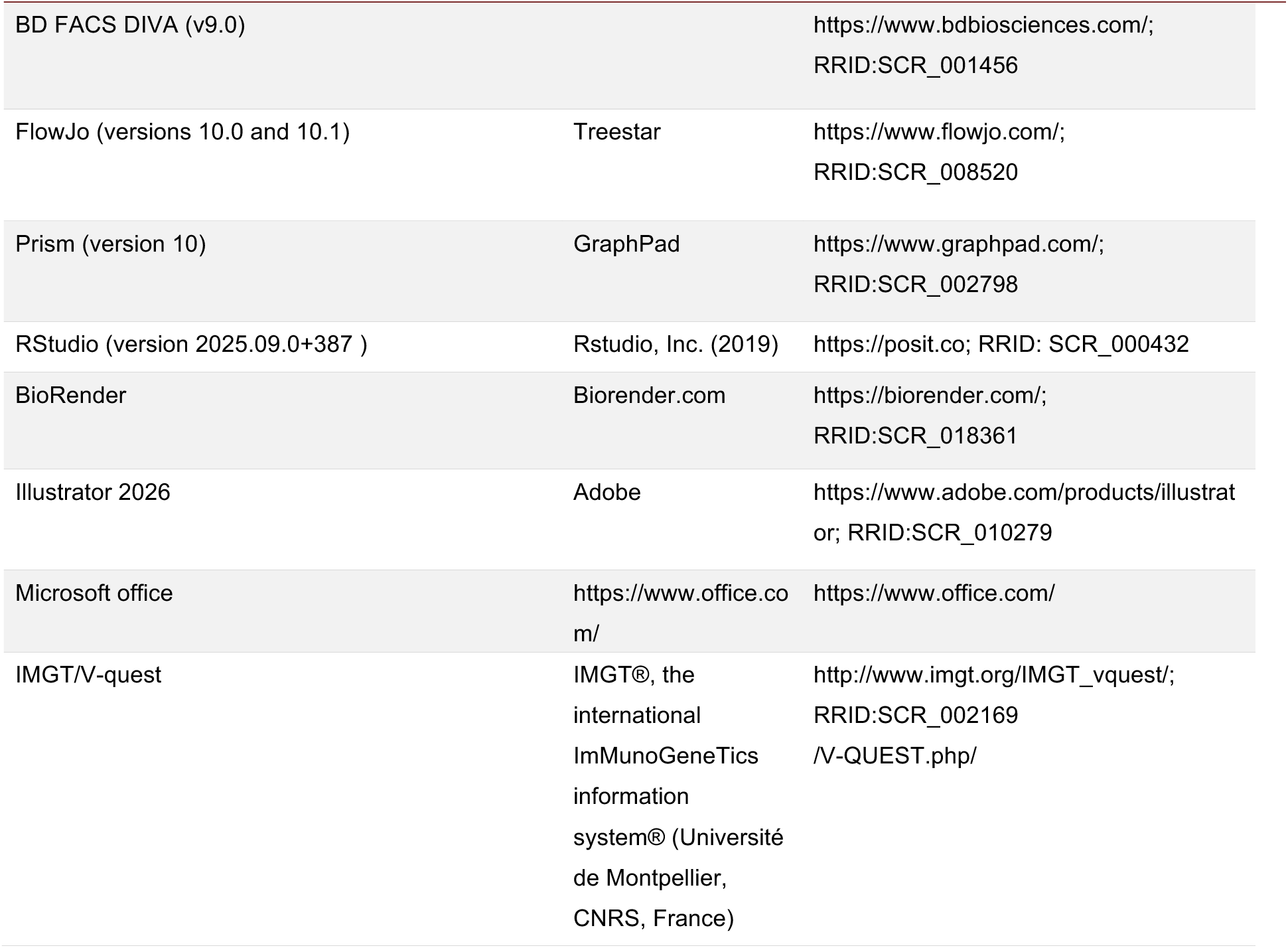

## SUPPLEMENTAL FIGURE LEGENDS

**Figure S1.**
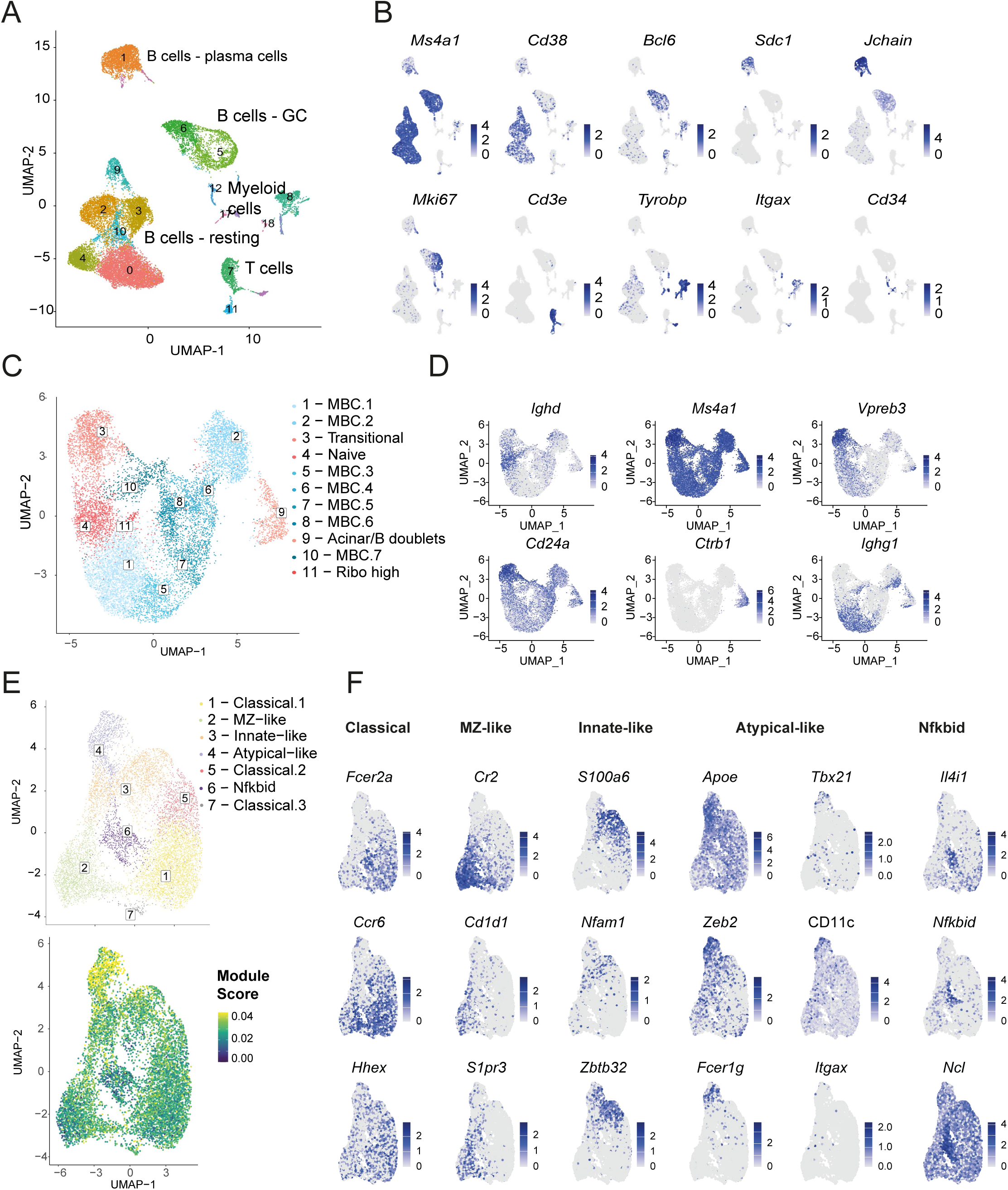
Overview of pre-subclustering UMAPs with final clustering results, related to Fig. 1. (A, B) UMAP of Harmony-integrated data from six experiments including the indicated MBC sorts (Fig. 1B) and sorts of other immune populations (A) with feature plots of cluster-defining marker genes (B). Clusters 0, 2, 3, 4, 9 and 10 correspond to resting B cells and were selected for further subclustering. (C, D) UMAP of the selected subset after subclustering (A), with feature plots showing defining marker genes. Transitional B cells (Cluster 3), naïve B cells (Cluster 4), Acinar/B doublets (Cluster 9) and ribo high cells (Cluster 1) were excluded prior to subclustering for the UMAP shown in Figure 1. (E) *Top:* UMAP and, *bottom:* feature plot showing the atypical MBC module score across all clusters. The score was calculated using Seurat’s AddModuleScore function based on the genes listed in Table S1. (F) Feature plots showing defining marker genes of clusters projected on UMAP shown in (E).

**Figure S2.**
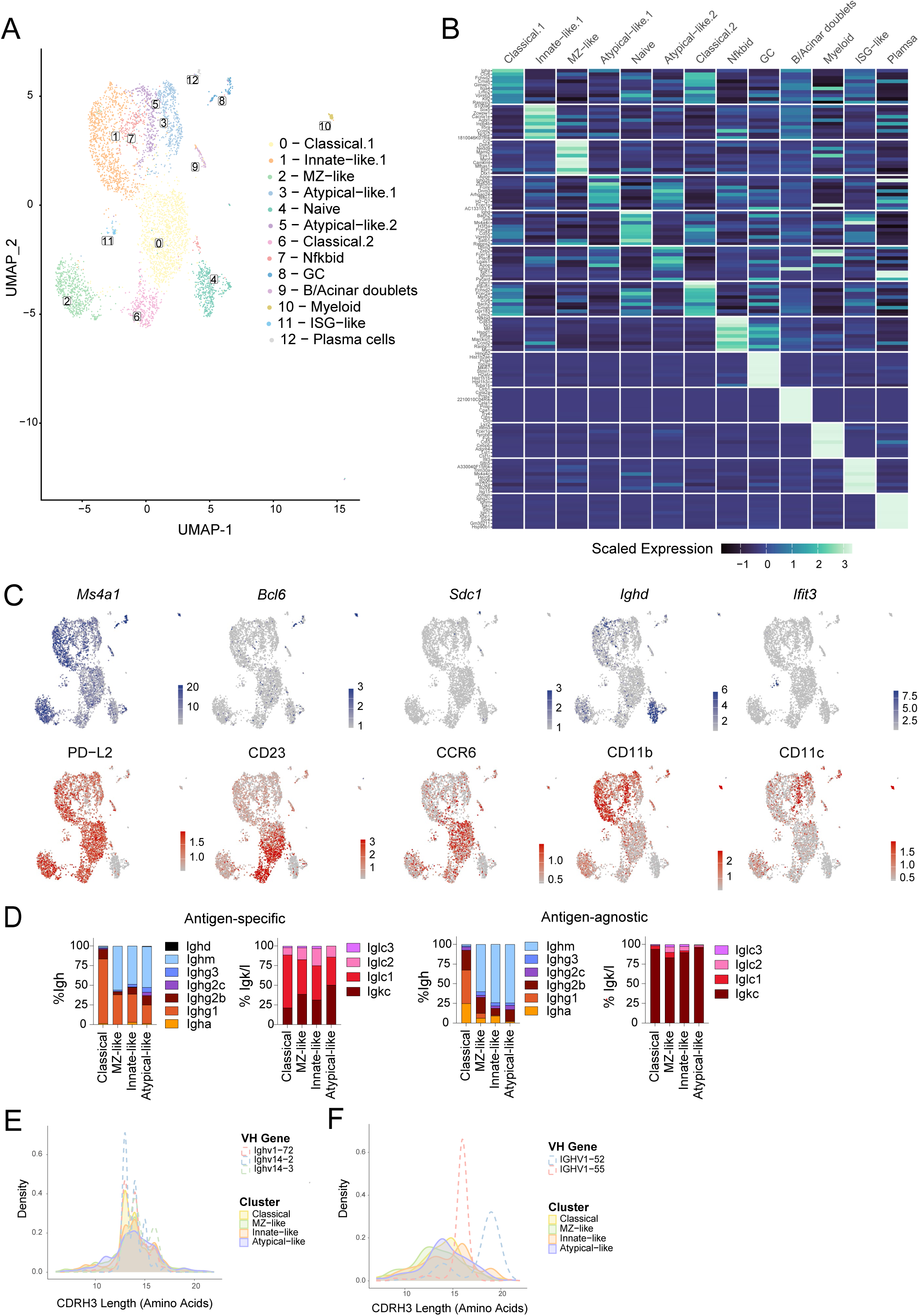
Additional clustering and CDRH3 length characterization, related to Figs. 1 and 2. (A) UMAP of a selected experiment from the Harmony-integrated dataset (Fig. 1) showing Louvain clusters at 0.8 resolution. (B) Heatmap showing the top 10 gene expression markers for each cluster. Color key: scaled mean expression. (C) Feature Plots showing expression of the indicated marker genes (*top*) and surface markers (*bottom*). (D) Frequency of Igh and Igk/l constant regions in antigen-specific (*left*) and antigen-agnostic cells (*right*) and per indicated MBC subset. (E, F) CDRH3 length of selected MBC clusters overlaid with the CDRH3 length distribution of (E) antigen-specific and (F) antigen-agnostic VH genes.

**Figure S3.**
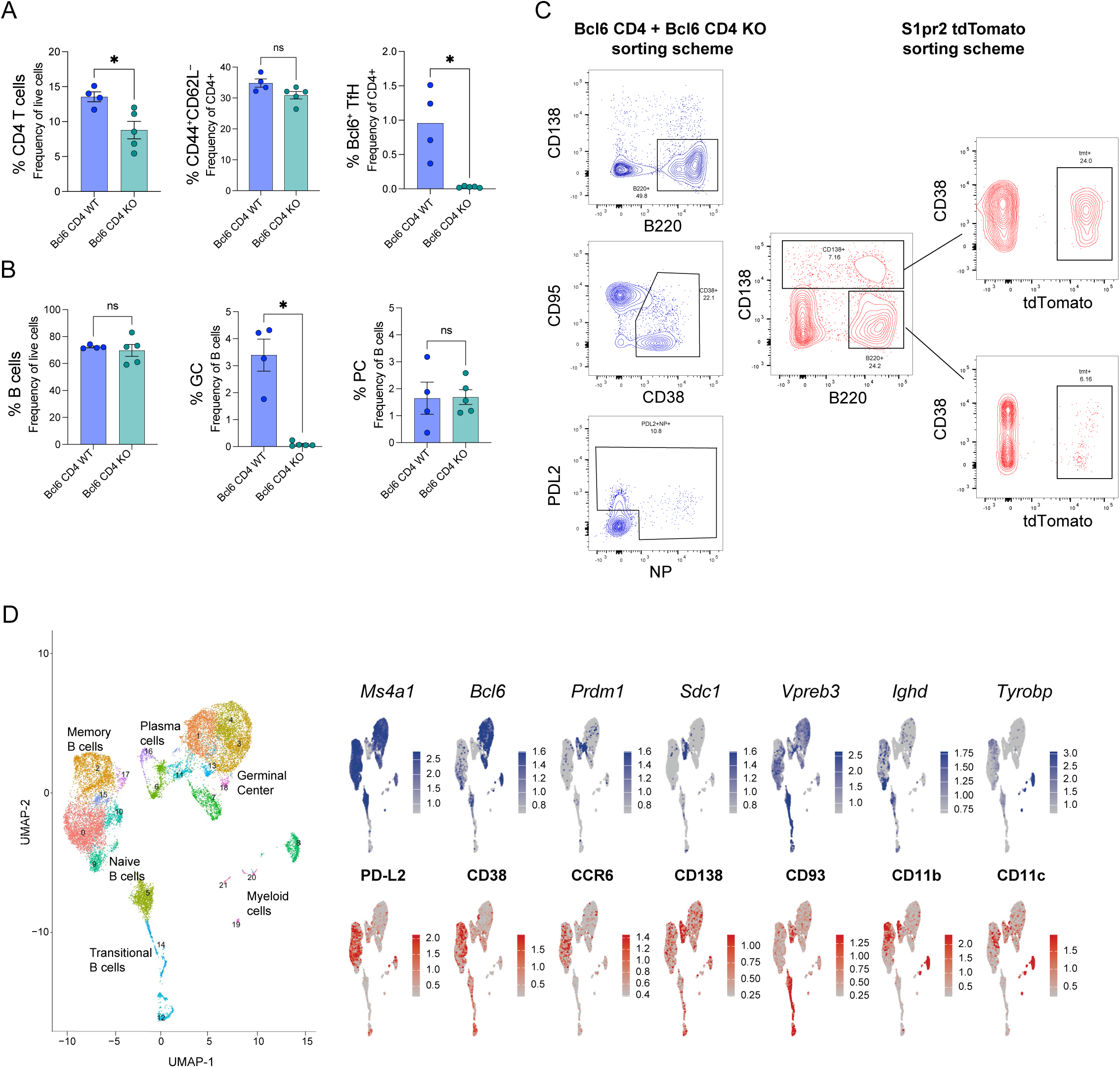
Establishing a workflow for single-cell sequencing comparisons between S1pr2 tdTomato and Bcl6 CD4 KO mice, related to Fig. 3. (A) Percentage of CD4^+^ T cells (*left*), activated CD44⁺CD62L⁻ T cells (*middle*), and Bcl6⁺ Tfh cells (*right*) in Bcl6 CD4 WT (blue) and Bcl6 CD4 KO (teal) mice. CD4^+^ T cells were gated as CD4⁺, activated T cells as CD4⁺CD25⁻CD44⁺CD62L⁻, and Tfh cells as CD4⁺CD25⁻CD44⁺CD62L⁻PD-1⁺CXCR5⁺Bcl6⁺. (B) Percentage of B cells (*left*), GC (*middle*) and PC (*right*) in Bcl6 CD4 WT (blue) and Bcl6 CD4 KO (teal) mice. B cells were gated as B220^+^, GC cells as B220^+^CD95^+^CD38^−^and PC as B220^lo^CD138^+^. (C) Sorting scheme for Bcl6 CD4 WT and Bcl6 CD4 KO mice (*left, blue*) and S1pr2 tdTomato mice (*right, red*). Cells from Bcl6 CD4 WT and Bcl6 CD4 KO mice were sorted as CD38^+^ (2000 cells/sample), PDL2^+^ and NP^+^. Cells from S1pr2 tdTomato mice were sorted as B220^+^tdTomato^+^ (8000 cells/sample) or CD138^+^ tdTomato^+^ (100 cells/sample). For WT, Bcl6 KO and S1pr2 tomato samples, three samples were sorted from mice that received primary NP-CGG immunization only and three samples from mice that received NP-CGG prime/boost immunization. (D) UMAP of Harmony-integrated data S1pr2 tdTomato, WT and Bcl6 KO mice sorted as described in panel (C) with corresponding feature plots of cluster-defining marker genes. Clusters 0, 2, 10, 15, and 17 were chosen for further subclustering for the UMAP shown in Fig. 3C.

**Figure S4.**
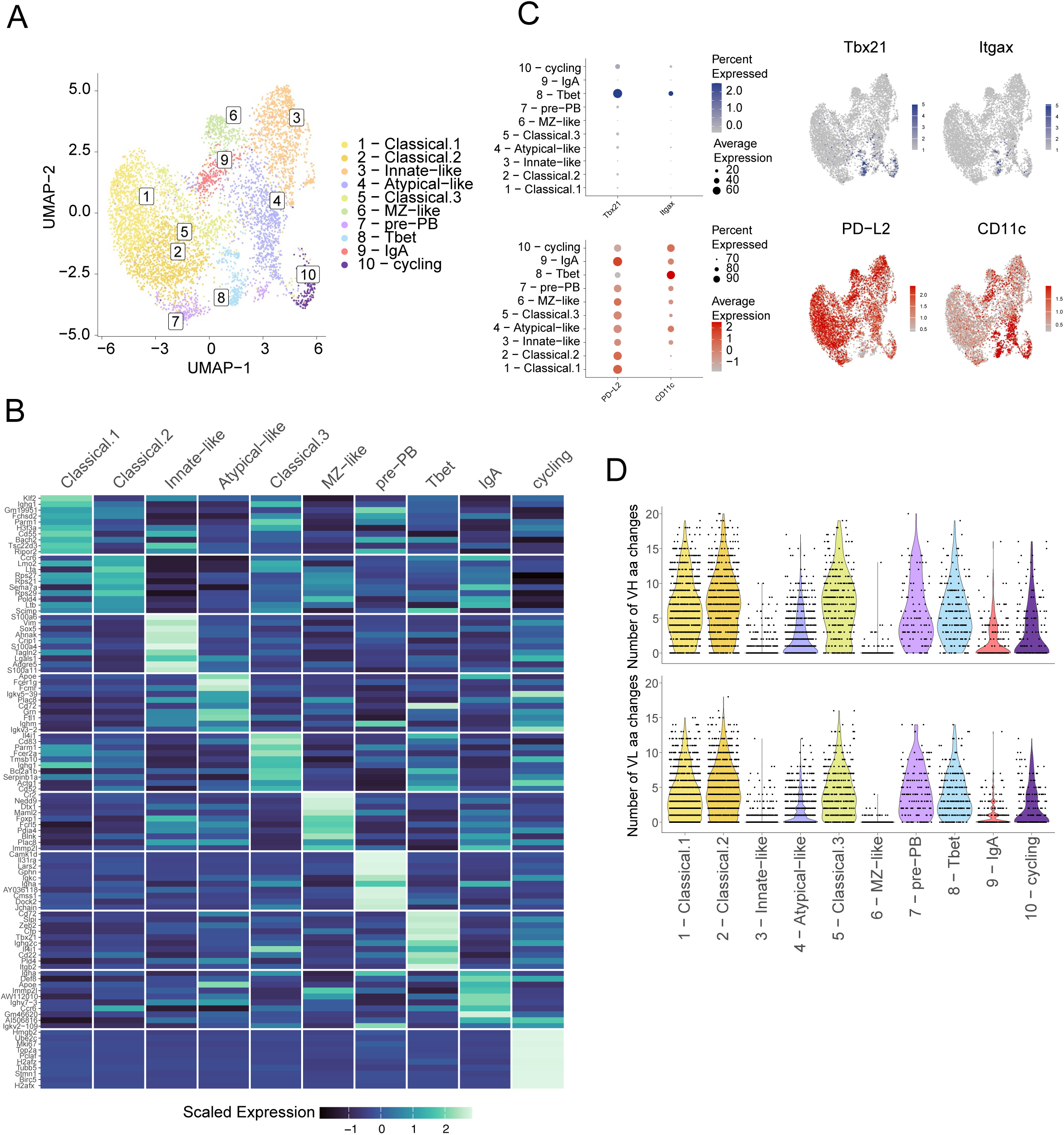
Cluster-defining markers and mutational profile across the S1pr2 tdTomato vs CD4 Bcl6 KO CITE-seq dataset, related to Fig. 3. (A) UMAP shown in Figure 3C for reference. (B) Heatmap showing the top 10 gene expression markers for each cluster from the S1pr2 tdTomato vs CD4 Bcl6 KO CITE-seq dataset. Color key: scaled mean expression. (C) Dotplots (*left*) and Feature Plots (*right*) showing expression of the indicated marker genes (*top*) and surface markers (*bottom*). Dot size represents the percentage of cells expressing each marker, and color intensity indicates scaled mean expression. Feature plots are overlaid on the UMAP embedding. (D) Violin plots showing the number of amino acid changes in VH (*top*) and VL (*bottom*) across clusters.

**Figure S5.**
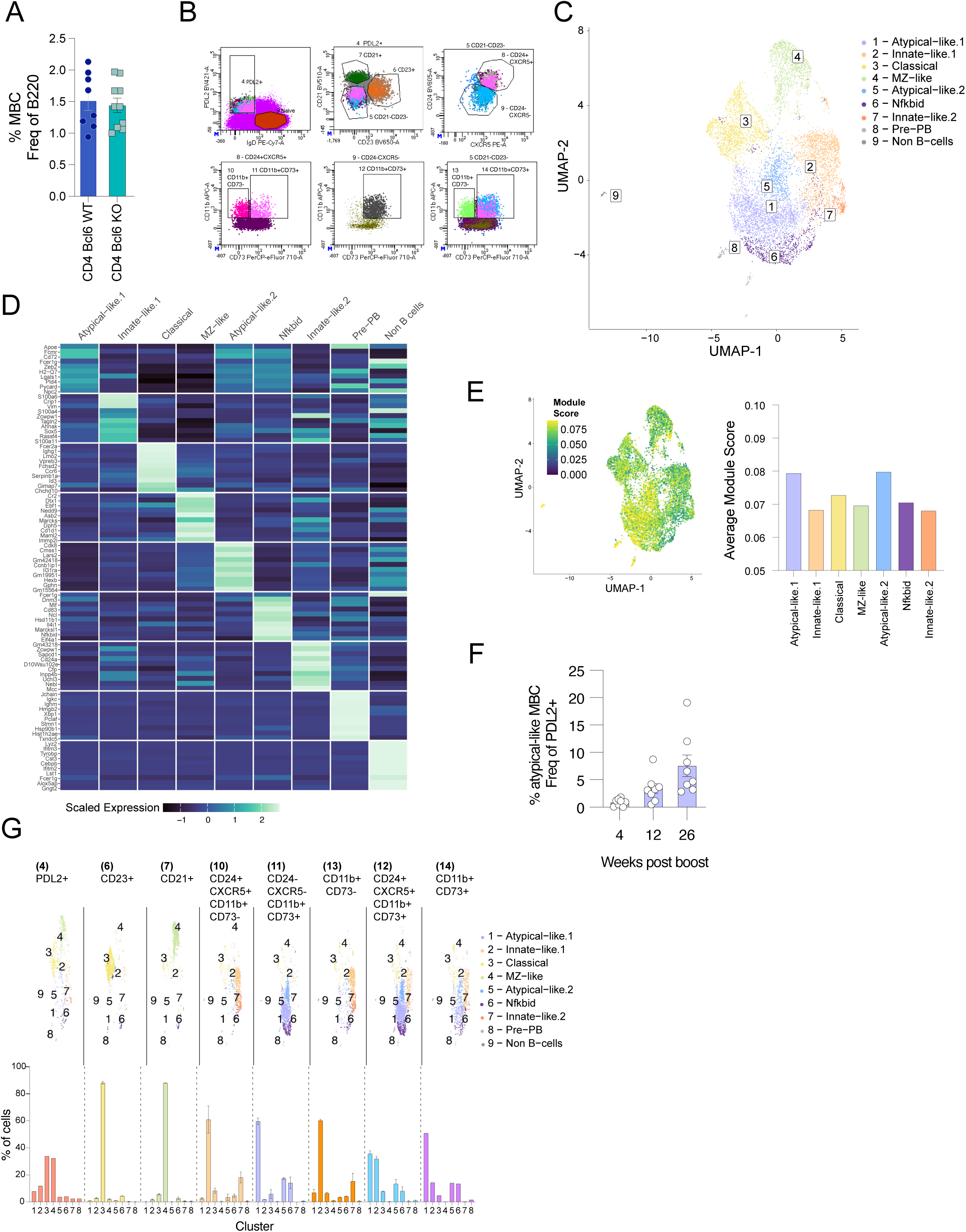
Single-cell sequencing demonstrates agreement between transcriptional profiles and surface marker–based cell populations, related to Fig. 4. (A) Frequency of total MBCs (B220^+^CD138^−^CD38^+^IgD^−^PDL2^+^) in Bcl6 CD4 WT vs Bcl6 CD4 KO chimeras. (B) Sorting scheme for single-cell experiment shown in Figure 4G&H. (C) Full UMAP shown in Figure 4G&H from cells sorted according to scheme in (A). (D) Heatmap showing the top 10 gene expression markers for each cluster from UMAP in (C). Color key: scaled mean expression. (E) *Left*: Feature Plot showing the atypical memory B cell modulescore across all clusters of UMAP shown in (B) *Right*: Bar plot showing the average module score per cluster (mean of per-cell module scores) across MBC clusters. (F) Frequency of atypical-like MBCs in naïve, immunized and boosted mice (Figure 1A) at different time points after boost. (G) *Top:* UMAP shown in (C) split by sorting condition. Numbering corresponds to gate assignments shown in (B). *Bottom*: Frequency of cells per cluster indicated for each sorting condition.

**Figure S6.**
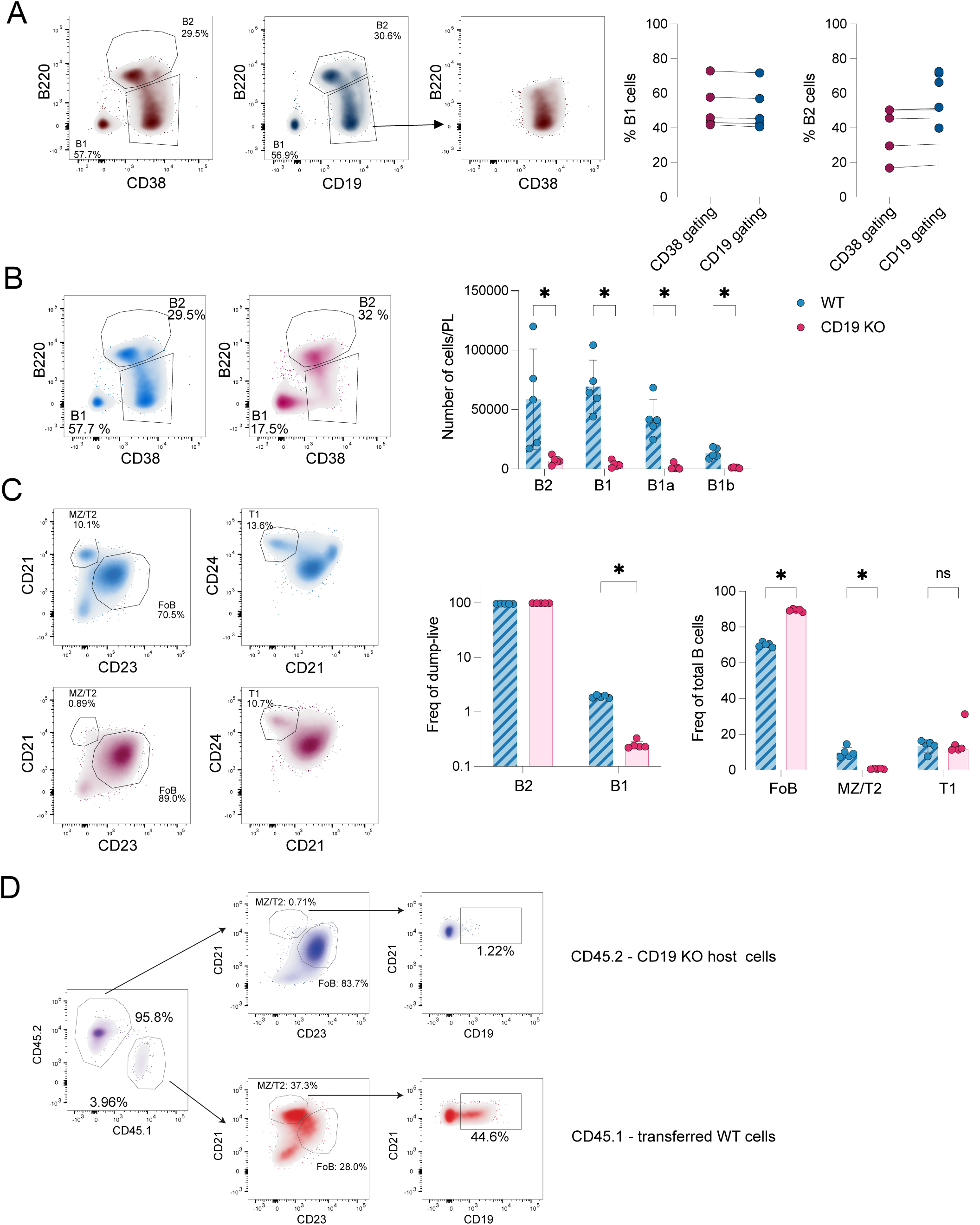
CD19 KO mice show significant impairment of the B1 and MZ compartments, related to Figure 5. (A) Alternative gating strategy to identify peritoneal B1 cells in CD19 KO animals. *Left*: Density plots showing cell populations obtained through CD38^+^B220^lo^ gating and through CD19^+^B220^lo^ gating. *Right*: Dot plots showing percentages of peritoneal B1 and B2 B cells by CD38 or CD19 gating. (B) *Left*: Contour plots of peritoneal B1 and B2 populations in WT (blue) and CD19 KO (pink) animals. *Right*: Number of cells per indicated cell populations in the PerCav of WT (blue) and CD19 KO (pink) animals. (C) *Left*: Contour plots of indicated splenic B cells populations in WT (blue) and CD19 KO (pink) animals. *Right*: Frequency of the indicated cell populations in the spleen of WT (blue) and CD19 KO (pink) animals. (D) Representative gating showing MZ reconstitution 10 weeks after B cell transfer. MZ and FoB gating is shown in CD45.2 host cells (blue) and CD45.1 adoptively transferred cells (red). Correct positioning of MZ cells is demonstrated by staining with CD19-PE which was injected i.v. 5 min prior to spleen harvest. Statistical significance was determined using an unpaired Mann-Whitney test with Holm-Sidak correction for multiple comparisons. In all bar plots, each symbol represents one mouse and mean ± SEM are marked. *p < 0.05, **p < 0.01, ***p < 0.001, and ****p < 0.0001.

**Figure S7.**
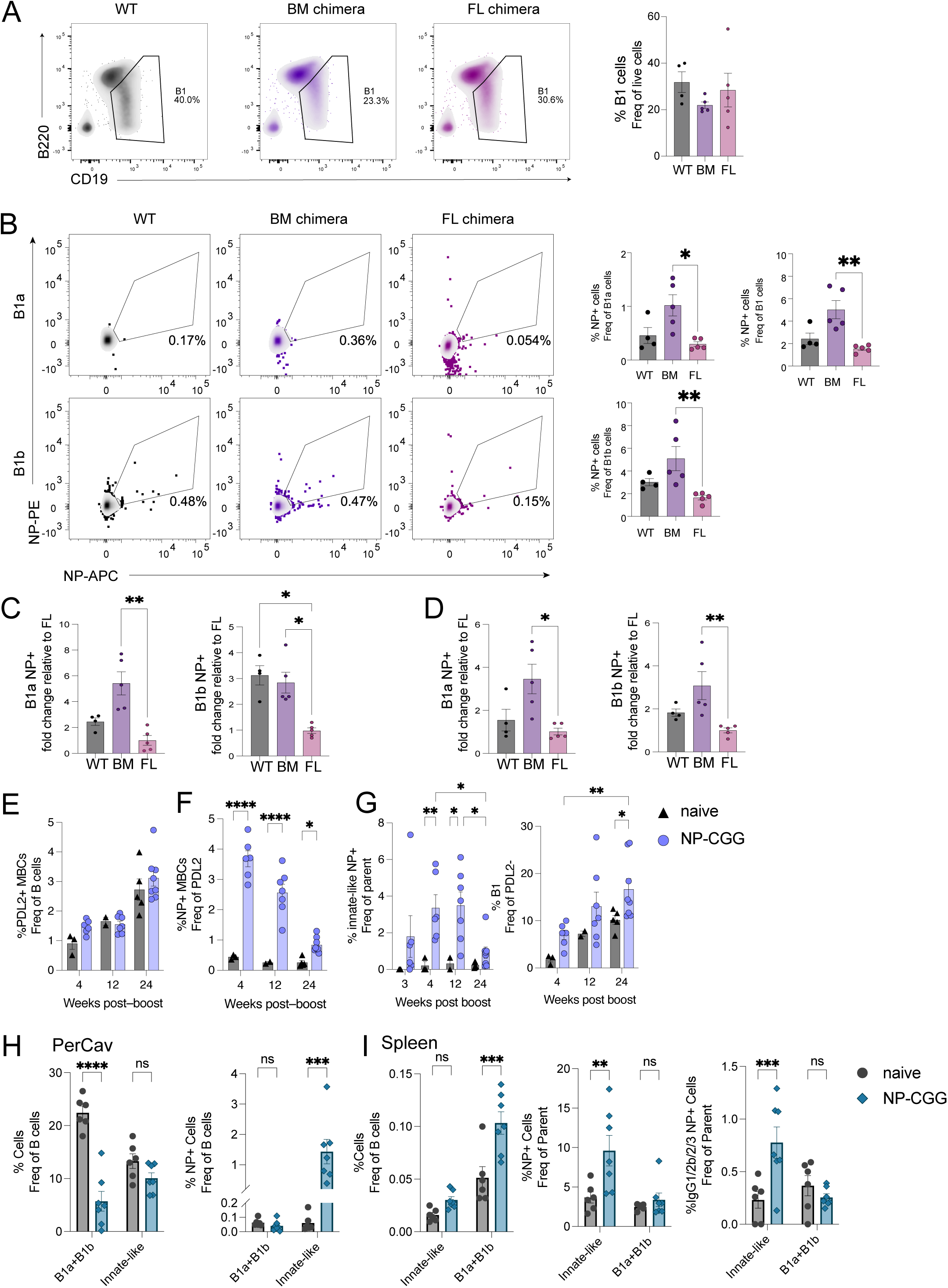
FL chimeric mice show reduced frequencies of antigen-specific cells, related to Figure 6. (A) Density plots (*left*) and quantifications (*right*) of total PerCav B1 cells in naïve WT (black), BM chimeric (purple), or FL chimeric (magenta) mice. (B) Density plots (*left*) and quantifications (*right*) of NP-specific cells in the indicated PerCav B1 populations in naïve WT, BM chimeric, or FL chimeric mice. (C) Fold change of antigen-specific peritoneal B1a (*left*) and B1b (*right*) in naïve WT, BM chimeric, and FL chimeric mice, relative to the mean of FL chimeric mice, set to 1. (D) Fold change of antigen-specific peritoneal B1a (*left*) and B1b (*right*) in immunized WT, BM chimeric and FL chimeric mice, relative to the mean of FL chimeric mice, set to 1. (E–I) Mice were immunized with 50 µg NP-CGG/AHG and boosted 4 weeks later with 50 µg NP-CGG/PBS. (E–G) Splenic B cell populations analyzed by flow cytometry at different weeks post-boost, as indicated on x-axis. (E) Quantification of total MBCs. (F) Frequency of total NP-specific MBCs. (G) Quantification of NP-specific innate-like MBCs (*left*) and B1 cells (*right*). (H) Frequencies of total B1 cells and innate-like MBCs (*left*) and NP-specific B1 and innate-like MBCs (*right*) in PerCav 6 weeks post-boost. (I) Bar plots depicting frequencies of total innate-like MBCs and B1 cells (*left*), NP-specific innate-like MBCs and B1 cells and (*middle*), NP-specific IgG1/2b/2c/3^+^ innate-like MBCs and B1 cells (*right*) in spleen 6 weeks post-boost. In all bar plots, each symbol represents one mouse and mean ± SEM are marked. *p < 0.05, **p < 0.01, ***p < 0.001, and ****p < 0.0001.

**Figure S8.**
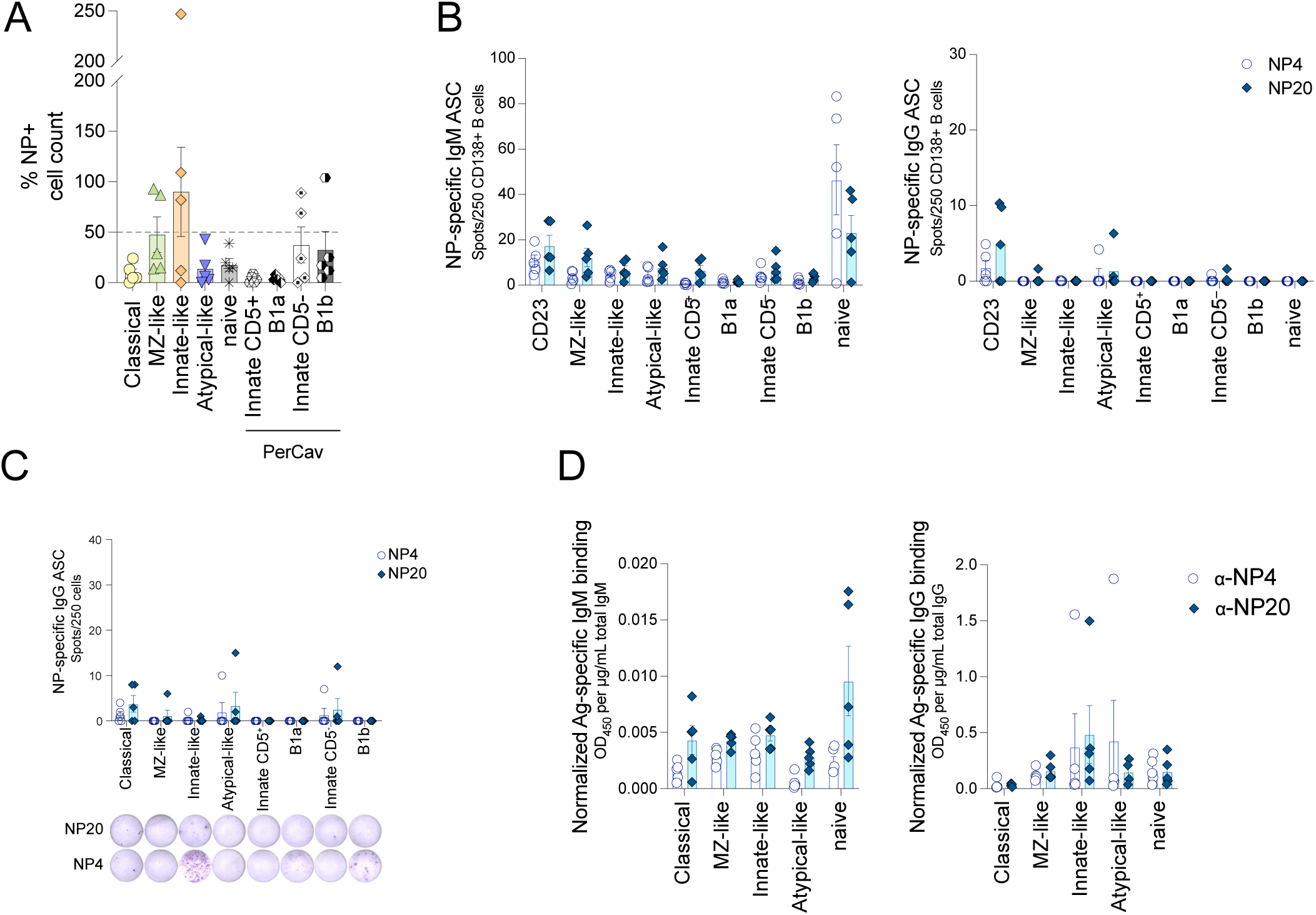
ASCs and OD450 normalized to CD138^+^ cells and antibody concentration, related to Fig. 7. (A) Quantification of antigen-specific NP^+^ cells by flow cytometry on Nojima culture day 4. (B) Antigen-specific IgM (*left*) and IgG (*right*) ASC counts normalized to 250 CD138^+^ cells. (C) Antigen-specific IgG ASC counts per 250 cells seeded at day 0 of Nojima culture. (D) OD₄₅₀ of antigen-specific IgM (*left*) and IgG (*right*), normalized to total IgM and IgG concentrations. In all bar plots, each symbol represents one mouse and mean ± SEM are marked.

## REFERENCES

Aiba, Yuichi, Kohei Kometani, Megumi Hamadate, Saya Moriyama, Asako Sakaue-Sawano, Michio Tomura, Hervé Luche, et al. 2010. ‘Preferential Localization of IgG Memory B Cells Adjacent to Contracted Germinal Centers’. Proceedings of the National Academy of Sciences 107(27): 12192–97. doi:10.1073/pnas.1005443107.

Allen, D., T. Simon, F. Sablitzky, K. Rajewsky, and A. Cumano. 1988. ‘Antibody Engineering for the Analysis of Affinity Maturation of an Anti-Hapten Response.’ The EMBO Journal 7(7): 1995–2001. doi:10.1002/j.1460-2075.1988.tb03038.x.

Alugupalli, Kishore R., John M. Leong, Robert T. Woodland, Masamichi Muramatsu, Tasuku Honjo, and Rachel M. Gerstein. 2004. ‘B1b Lymphocytes Confer T Cell-Independent Long-Lasting Immunity’. Immunity 21(3): 379–90. doi:10.1016/j.immuni.2004.06.019.

Amano, Masahiko, Nicole Baumgarth, Michael D. Dick, Laurent Brossay, Mitchell Kronenberg, Lee A. Herzenberg, and Samuel Strober. 1998. ‘CD1 Expression Defines Subsets of Follicular and Marginal Zone B Cells in the Spleen: Β2-Microglobulin-Dependent and Independent Forms’. Journal of Immunology 161(4): 1710–17.

Arnold, Carrie N., Elaine Pirie, Pia Dosenovic, Gerald M. McInerney, Yu Xia, Nathaniel Wang, Xiaohong Li, et al. 2012. ‘A Forward Genetic Screen Reveals Roles for Nfkbid, Zeb1, and Ruvbl2 in Humoral Immunity’. Proceedings of the National Academy of Sciences of the United States of America 109(31): 12286–93. doi:10.1073/pnas.1209134109.

Arnon, Tal I., Robert M. Horton, Irina L. Grigorova, and Jason G. Cyster. 2013. ‘Visualization of Splenic Marginal Zone B Cell Shuttling and Follicular B Cell Egress’. Nature 493(7434): 684–88. doi:10.1038/nature11738.

Baumgarth, Nicole. 2011. ‘The Double Life of a B-1 Cell: Self-Reactivity Selects for Protective Effector Functions’. Nature Reviews Immunology 11(1): 34–46. doi:10.1038/nri2901.

Baumgarth, Nicole. 2016. ‘B-1 Cell Heterogeneity and the Regulation of Natural and Antigen-Induced IgM Production’. Frontiers in Immunology 7: 324. doi:10.3389/fimmu.2016.00324.

Baumgarth, Nicole. 2017. ‘A Hard(y) Look at B-1 Cell Development and Function*’. Journal of immunology (Baltimore, Md.: 1950) 199(10): 3387–94. doi:10.4049/jimmunol.1700943.

Baumgarth, Nicole, Ometa C. Herman, Gina C. Jager, Lorena E. Brown, Leonore A. Herzenberg, and Jianzhu Chen. 2000. ‘B-1 and B-2 Cell–Derived Immunoglobulin M Antibodies Are Nonredundant Components of the Protective Response to Influenza Virus Infection’. The Journal of Experimental Medicine 192(2): 271–80. doi:10.1084/jem.192.2.271.

Bod, Lloyd, Laetitia Douguet, Cédric Auffray, Renée Lengagne, Fériel Bekkat, Elena Rondeau, Valérie Molinier-Frenkel, et al. 2018. ‘IL-4-Induced Gene 1: A Negative Immune Checkpoint Controlling B Cell Differentiation and Activation’. Journal of Immunology 200(3): 1027–38. doi:10.4049/jimmunol.1601609.

Boes, M. 2000. ‘Role of Natural and Immune IgM Antibodies in Immune Responses’. Molecular Immunology 37(18): 1141–49. doi:10.1016/s0161-5890(01)00025-6.

Boes, Marianne, Andrey P. Prodeus, Tara Schmidt, Michael C. Carroll, and Jianzhu Chen. 1998. ‘A Critical Role of Natural Immunoglobulin M in Immediate Defense Against Systemic Bacterial Infection’. The Journal of Experimental Medicine 188(12): 2381–86. doi:10.1084/jem.188.12.2381.

Callahan, Derrick, Shuchi Smita, Stephen Joachim, Kenneth Hoehn, Steven Kleinstein, Florian Weisel, Maria Chikina, and Mark Shlomchik. 2024. ‘Memory B Cell Subsets Have Divergent Developmental Origins That Are Coupled to Distinctive Imprinted Epigenetic States’. Nature immunology 25(3): 562–75. doi:10.1038/s41590-023-01721-9.

Casali, P., and E. W. Schettino. 1996. ‘Structure and Function of Natural Antibodies’. Current Topics in Microbiology and Immunology 210: 167–79. doi:10.1007/978-3-642-85226-8_17.

Choi, Youn Soo, Jacquelyn A. Dieter, Kristina Rothaeusler, Zheng Luo, and Nicole Baumgarth. 2012. ‘B-1 Cells in the Bone Marrow Are a Significant Source of Natural IgM’. European journal of immunology 42(1): 120–29. doi:10.1002/eji.201141890.

Cole, Leah E., Yang Yang, Karen L. Elkins, Ellen T. Fernandez, Nilofer Qureshi, Mark J. Shlomchik, Leonard A. Herzenberg, Leonore A. Herzenberg, and Stefanie N. Vogel. 2009. ‘Antigen-Specific B-1a Antibodies Induced by *Francisella Tularensis* LPS Provide Long-Term Protection against *F. Tularensis* LVS Challenge’. Proceedings of the National Academy of Sciences 106(11): 4343–48. doi:10.1073/pnas.0813411106.

Cooper, Lucy, Hui Xu, Jack Polmear, Liam Kealy, Christopher Szeto, Ee Shan Pang, Mansi Gupta, et al. 2024. ‘Type I Interferons Induce an Epigenetically Distinct Memory B Cell Subset in Chronic Viral Infection’. Immunity 57(5): 1037–1055.e6. doi:10.1016/j.immuni.2024.03.016.

Cumano, A., and K. Rajewsky. 1986. ‘Clonal Recruitment and Somatic Mutation in the Generation of Immunological Memory to the Hapten NP’. The EMBO journal 5(10): 2459–68. doi:10.1002/j.1460-2075.1986.tb04522.x.

Cunningham, Adam F., Adriana Flores-Langarica, Saeeda Bobat, Carmen C. Dominguez Medina, Charlotte N. L. Cook, Ewan A. Ross, Constantino Lopez-Macias, and Ian R. Henderson. 2014. ‘B1b Cells Recognize Protective Antigens after Natural Infection and Vaccination’. Frontiers in Immunology 5: 535. doi:10.3389/fimmu.2014.00535.

Dai, Dai, Shuangshuang Gu, Xiaxia Han, Huihua Ding, Yang Jiang, Xiaoou Zhang, Chao Yao, et al. 2024. ‘The Transcription Factor ZEB2 Drives the Formation of Age-Associated B Cells’. Science 383(6681): 413–21. doi:10.1126/science.adf8531.

Du, Samuel W, Tanvi Arkatkar, Fahd Al Qureshah, Holly M Jacobs, Christopher D Thouvenel, Kristy Chiang, Andrea D Largent, et al. 2019. ‘Functional Characterization of CD11c+ Age-Associated B Cells as Memory B Cells’. The Journal of Immunology 203(11): 2817–26. doi:10.4049/jimmunol.1900404.

Duan, Lihui, Dan Liu, Hsin Chen, Michelle A. Mintz, Marissa Y. Chou, Dmitri I. Kotov, Ying Xu, et al. 2021. ‘Follicular Dendritic Cells Restrict Interleukin-4 Availability in Germinal Centers and Foster Memory B Cell Generation’. Immunity 54(10): 2256–2272.e6. doi:10.1016/j.immuni.2021.08.028.

Ehrenstein, Michael R., and Clare A. Notley. 2010. ‘The Importance of Natural IgM: Scavenger, Protector and Regulator’. Nature Reviews Immunology 10(11): 778–86. doi:10.1038/nri2849.

Elgueta, Raul, Ellen Marks, Elizabeth Nowak, Shinelle Menezes, Micah Benson, Vanitha S. Raman, Carla Ortiz, et al. 2015. ‘CCR6-Dependent Positioning of Memory B Cells Is Essential for Their Ability to Mount a Recall Response to Antigen’. Journal of immunology (Baltimore, Md.: 1950) 194(2): 505–13. doi:10.4049/jimmunol.1401553.

Fremerey, Julia, Pavel Morozov, Cindy Meyer, Aitor Garzia, Marianna Teplova, Thomas Tuschl, and Arndt Borkhardt. 2016. ‘Nucleolin Controls Ribosome Biogenesis through Its RNA-Binding Properties’. Blood 128(22): 5056–5056. doi:10.1182/blood.V128.22.5056.5056.

Gao, Xin, Qian Shen, Jonathan A. Roco, Becan Dalton, Katie Frith, C. Mee Ling Munier, Fiona D. Ballard, et al. 2024. ‘Zeb2 Drives the Formation of CD11c^+^ Atypical B Cells to Sustain Germinal Centers That Control Persistent Infection’. Science Immunology 9(93): eadj4748. doi:10.1126/sciimmunol.adj4748.

Green, Jesse A., Kazuhiro Suzuki, Bryan Cho, L. David Willison, Daniel Palmer, Christopher D.C. Allen, Timothy H. Schmidt, et al. 2011. ‘The Sphingosine 1-Phosphate Receptor S1P2 Maintains Germinal Center B Cell Homeostasis and Promotes Niche Confinement’. Nature immunology 12(7): 672–80. doi:10.1038/ni.2047.

Gregoire, Claude, Lionel Spinelli, Sergio Villazala-Merino, Laurine Gil, María Pía Holgado, Myriam Moussa, Chuang Dong, et al. 2022. ‘Viral Infection Engenders Bona Fide and Bystander Subsets of Lung-Resident Memory B Cells through a Permissive Mechanism’. Immunity 55(7): 1216–1233.e9. doi:10.1016/j.immuni.2022.06.002.

Guo, Benchang, Alexander V. Ludlow, Angela S. Brightwell, and Thomas L. Rothstein. 2019. ‘Impairment of PD-L2 Positive B1a Cells Enhances Susceptibility to Sepsis in RasGRP1-Deficient Mice’. Cellular Immunology 346: 103993. doi:10.1016/j.cellimm.2019.103993.

Haas, Karen M., Jonathan C. Poe, Douglas A. Steeber, and Thomas F. Tedder. 2005. ‘B-1a and B-1b Cells Exhibit Distinct Developmental Requirements and Have Unique Functional Roles in Innate and Adaptive Immunity to S. Pneumoniae’. Immunity 23(1): 7–18. doi:10.1016/j.immuni.2005.04.011.

Haase, Paul, Simon Schäfer, Roman G. Gerlach, Thomas H. Winkler, and David Voehringer. 2022. ‘B Cell Fate Mapping Reveals Their Contribution to the Memory Immune Response against Helminths’. Frontiers in Immunology 13: 1016142. doi:10.3389/fimmu.2022.1016142.

Hanakahi, L. A., L. A. Dempsey, M. J. Li, and N. Maizels. 1997. ‘Nucleolin Is One Component of the B Cell-Specific Transcription Factor and Switch Region Binding Protein, LR1’. Proceedings of the National Academy of Sciences of the United States of America 94(8): 3605–10. doi:10.1073/pnas.94.8.3605.

Hao, Yi, Patrick O’Neill, Martin S. Naradikian, Jean L. Scholz, and Michael P. Cancro. 2011. ‘A B-Cell Subset Uniquely Responsive to Innate Stimuli Accumulates in Aged Mice’. Blood 118(5): 1294–1304. doi:10.1182/blood-2011-01-330530.

Hao, Yuhan, Stephanie Hao, Erica Andersen-Nissen, William M. Mauck, Shiwei Zheng, Andrew Butler, Maddie J. Lee, et al. 2021. ‘Integrated Analysis of Multimodal Single-Cell Data’. Cell 184(13): 3573–3587.e29. doi:10.1016/j.cell.2021.04.048.

Hayakawa, Kyoko, Anthony M. Formica, Joni Brill-Dashoff, Susan A. Shinton, Daiju Ichikawa, Yan Zhou, Herbert C. Morse, and Richard R. Hardy. 2016. ‘Early Generated B1 B Cells with Restricted BCRs Become Chronic Lymphocytic Leukemia with Continued C-Myc and Low Bmf Expression’. The Journal of Experimental Medicine 213(13): 3007–24. doi:10.1084/jem.20160712.

Hendricks, Jacobus, Nicolaas A. Bos, and Frans G.M. Kroese. 2018. ‘Heterogeneity of Memory Marginal Zone B Cells’. Critical reviews in immunology 38(2): 145–58. doi:10.1615/CritRevImmunol.2018024985.

Hsu, Mei-Chi, Kai-Michael Toellner, Carola G. Vinuesa, and Ian C. M. MacLennan. 2006. ‘B Cell Clones That Sustain Long-Term Plasmablast Growth in T-Independent Extrafollicular Antibody Responses’. Proceedings of the National Academy of Sciences 103(15): 5905–10. doi:10.1073/pnas.0601502103.

Jash, Arijita, Yinan Wang, Florian J. Weisel, Christopher D. Scharer, Jeremy M. Boss, Mark J. Shlomchik, and Deepta Bhattacharya. 2016. ‘ZBTB32 Restricts the Duration of Memory B Cell Recall Responses’. Journal of immunology (Baltimore, Md.: 1950) 197(4): 1159–68. doi:10.4049/jimmunol.1600882.

Jash, Arijita, You W. Zhou, Diana K. Gerardo, Tyler J. Ripperger, Bijal A. Parikh, Sytse Piersma, Deepa R. Jamwal, et al. 2019. ‘ZBTB32 Restrains Antibody Responses to Murine Cytomegalovirus Infections, but Not Other Repetitive Challenges’. Scientific Reports 9(1): 15257. doi:10.1038/s41598-019-51860-z.

Johnson, John L, Rebecca L Rosenthal, James J Knox, Arpita Myles, Martin S Naradikian, Joanna Madej, Mariya Kostiv, et al. 2020. ‘The Transcription Factor T-Bet Resolves Memory B Cell Subsets with Distinct Tissue Distributions and Antibody Specificities in Mice and Humans’. Immunity 52(5): 842–855.e6. doi:10.1016/j.immuni.2020.03.020.

Juchem, Kathryn W., Anshu P. Gounder, Jian Ping Gao, Elise Seccareccia, Narayana Yeddula, Nicholas J. Huffmaster, Alexandra Côté-Martin, et al. 2021. ‘NFAM1 Promotes Pro-Inflammatory Cytokine Production in Mouse and Human Monocytes’. Frontiers in Immunology 12: 773445. doi:10.3389/fimmu.2021.773445.

Kaji, Tomohiro, Akiko Ishige, Masaki Hikida, Junko Taka, Atsushi Hijikata, Masato Kubo, Takeshi Nagashima, et al. 2012. ‘Distinct Cellular Pathways Select Germline-Encoded and Somatically Mutated Antibodies into Immunological Memory’. The Journal of Experimental Medicine 209(11): 2079–97. doi:10.1084/jem.20120127.

Kajon, Adriana E., Corrie C. Brown, and Katherine R. Spindler. 1998. ‘Distribution of Mouse Adenovirus Type 1 in Intraperitoneally and Intranasally Infected Adult Outbred Mice’. Journal of Virology 72(2): 1219–23. doi:10.1128/jvi.72.2.1219-1223.1998.

Khoenkhoen, Sharesta, Monika Ádori, Darío Solís-Sayago, Juliette Soulier, Jamie Russell, Bruce Beutler, Gabriel K. Pedersen, and Gunilla B. Karlsson Hedestam. 2022. ‘IκBNS Expression in B Cells Is Dispensable for IgG Responses to T Cell-Dependent Antigens’. Frontiers in Immunology 13: 1000755. doi:10.3389/fimmu.2022.1000755.

Knox, James J., Arpita Myles, and Michael P. Cancro. 2019. ‘T-bet^+^ Memory B Cells: Generation, Function, and Fate’. Immunological Reviews 288(1): 149–60. doi:10.1111/imr.12736.

Korsunsky, Ilya, Nghia Millard, Jean Fan, Kamil Slowikowski, Fan Zhang, Kevin Wei, Yuriy Baglaenko, et al. 2019. ‘Fast, Sensitive and Accurate Integration of Single-Cell Data with Harmony’. Nature Methods 16(12): 1289–96. doi:10.1038/s41592-019-0619-0.

Kozlyuk, Natalia, Andrew J. Monteith, Velia Garcia, Steven M. Damo, Eric P. Skaar, and Walter J. Chazin. 2019. ‘S100 Proteins in the Innate Immune Response to Pathogens’. Methods in molecular biology (Clifton, N.J.) 1929: 275–90. doi:10.1007/978-1-4939-9030-6_18.

Laidlaw, Brian J., Lihui Duan, Ying Xu, Sara E. Vazquez, and Jason G. Cyster. 2020. ‘The Transcription Factor Hhex Cooperates with the Corepressor Tle3 to Promote Memory B Cell Development’. Nature immunology 21(9): 1082–93. doi:10.1038/s41590-020-0713-6.

Leeman-Neill, Rebecca J., Junghyun Lim, and Uttiya Basu. 2018. ‘The Common Key to Class-Switch Recombination and Somatic Hypermutation: Discovery of AID and Its Role in Antibody Gene Diversification’. Journal of immunology (Baltimore, Md.: 1950) 201(9): 2527–29. doi:10.4049/jimmunol.1801246.

Li, Jiawei, Yi Zhang, Cheng Yang, and Ruiming Rong. 2020. ‘Discrepant mRNA and Protein Expression in Immune Cells’. Current Genomics 21(8): 560–63. doi:10.2174/1389202921999200716103758.

Liu, Dan, Benjamin Y. Winer, Marissa Y. Chou, Hanson Tam, Ying Xu, Jinping An, James M. Gardner, and Jason G. Cyster. 2024. ‘Dynamic Encounters with Red Blood Cells Trigger Splenic Marginal Zone B Cell Retention and Function’. Nature Immunology 25(1): 142–54. doi:10.1038/s41590-023-01690-z.

Luo, Yao, Jing Wang, Kairui Li, Mingxia Li, Shasha Xu, Xingjie Liu, Zhiwei Zhang, et al. 2022. ‘Single-Cell Genomics Identifies Distinct B1 Cell Developmental Pathways and Reveals Aging-Related Changes in the B-Cell Receptor Repertoire’. Cell & Bioscience 12: 57. doi:10.1186/s13578-022-00795-6.

Madisen, Linda, Theresa A Zwingman, Susan M Sunkin, Seung Wook Oh, Hatim A Zariwala, Hong Gu, Lydia L Ng, et al. 2010. ‘A Robust and High-Throughput Cre Reporting and Characterization System for the Whole Mouse Brain’. Nature Neuroscience 13(1): 133–40. doi:10.1038/nn.2467.

Marshall, Jennifer L, Adriana Flores-Langarica, Robert A Kingsley, Jessica R Hitchcock, Ewan A Ross, Constantino López-Macías, Jeremy Lakey, et al. 2012. ‘The Capsular Polysaccharide Vi from *Salmonella* Typhi Is a B1b Antigen’. The Journal of Immunology 189(12): 5527–32. doi:10.4049/jimmunol.1103166.

Mattos, Matheus Silvério, Sofie Vandendriessche, Ari Waisman, and Pedro Elias Marques. 2024. ‘The Immunology of B-1 Cells: From Development to Aging’. Immunity & Ageing: I & A 21: 54. doi:10.1186/s12979-024-00455-y.

Mesin, Luka, Ariën Schiepers, Jonatan Ersching, Alexandru Barbulescu, Cecília B. Cavazzoni, Alessandro Angelini, Takaharu Okada, Tomohiro Kurosaki, and Gabriel D. Victora. 2020. ‘Restricted Clonality and Limited Germinal Center Reentry Characterize Memory B Cell Reactivation by Boosting’. Cell 180(1): 92–106.e11. doi:10.1016/j.cell.2019.11.032.

Moore, Martin L., Corrie C. Brown, and Katherine R. Spindler. 2003. ‘T Cells Cause Acute Immunopathology and Are Required for Long-Term Survival in Mouse Adenovirus Type 1-Induced Encephalomyelitis’. Journal of Virology 77(18): 10060–70. doi:10.1128/JVI.77.18.10060-10070.2003.

Nojima, Takuya, Kei Haniuda, Tatsuya Moutai, Moeko Matsudaira, Sho Mizokawa, Ikuo Shiratori, Takachika Azuma, and Daisuke Kitamura. 2011. ‘In-Vitro Derived Germinal Centre B Cells Differentially Generate Memory B or Plasma Cells in Vivo’. Nature Communications 2: 465. doi:10.1038/ncomms1475.

Onodera, Taishi, Yoshimasa Takahashi, Yusuke Yokoi, Manabu Ato, Yuichi Kodama, Satoshi Hachimura, Tomohiro Kurosaki, and Kazuo Kobayashi. 2012. ‘Memory B Cells in the Lung Participate in Protective Humoral Immune Responses to Pulmonary Influenza Virus Reinfection’. Proceedings of the National Academy of Sciences of the United States of America 109(7): 2485–90. doi:10.1073/pnas.1115369109.

Palma, Joanna, Beata Tokarz-Deptuła, Jakub Deptuła, and Wiesław Deptuła. 2018. ‘Natural Antibodies – Facts Known and Unknown’. Central-European Journal of Immunology 43(4): 466–75. doi:10.5114/ceji.2018.81354.

Pape, Kathryn A., Justin J. Taylor, Robert W. Maul, Patricia J. Gearhart, and Marc K. Jenkins. 2011. ‘Different B Cell Populations Mediate Early and Late Memory During an Endogenous Immune Response’. Science 331(6021): 1203–7. doi:10.1126/science.1201730.

Purtha, Whitney E., Thomas F. Tedder, Syd Johnson, Deepta Bhattacharya, and Michael S. Diamond. 2011. ‘Memory B Cells, but Not Long-Lived Plasma Cells, Possess Antigen Specificities for Viral Escape Mutants’. Journal of Experimental Medicine 208(13): 2599–2606. doi:10.1084/jem.20110740.

Reth, M., G. J. Hämmerling, and K. Rajewsky. 1978. ‘Analysis of the Repertoire of anti-NP Antibodies in C57BL/6 Mice by Cell Fusion. I. Characterization of Antibody Families in the Primary and Hyperimmune Response’. European Journal of Immunology 8(6): 393–400. doi:10.1002/eji.1830080605.

Ridderstad, Anna, and David M Tarlinton. 1998. ‘Kinetics of Establishing the Memory B Cell Population as Revealed by CD38 Expression’. The Journal of Immunology 160(10): 4688–95. doi:10.4049/jimmunol.160.10.4688.

Riedel, René, Richard Addo, Marta Ferreira-Gomes, Gitta Anne Heinz, Frederik Heinrich, Jannis Kummer, Victor Greiff, et al. 2020. ‘Discrete Populations of Isotype-Switched Memory B Lymphocytes Are Maintained in Murine Spleen and Bone Marrow’. Nature Communications 11(1): 2570. doi:10.1038/s41467-020-16464-6.

Risley, Christopher A, Christopher D Scharer, Jeremy M Boss, and Frances E Lund. 2021. ‘T-Bet Expression Marks a Transcriptionally and Functionally Distinct Population of Memory B Cells’. The Journal of Immunology 206(1_Supplement): 114.18. doi:10.4049/jimmunol.206.Supp.114.18.

Risley, Christopher A., Michael D. Schultz, S. Rameeza Allie, Shanrun Liu, Jessica N. Peel, Anoma Nellore, Christopher F. Fucile, et al. 2025. ‘Transcription Factor T-Bet Regulates the Maintenance and Differentiation Potential of Lymph Node and Lung Effector Memory B Cell Subsets’. Immunity 58(7): 1706–1724.e6. doi:10.1016/j.immuni.2025.05.021.

Robbiani, Davide F., Anne Bothmer, Elsa Callen, Bernardo Reina-San-Martin, Yair Dorsett, Simone Difilippantonio, Daniel J. Bolland, et al. 2008. ‘AID Is Required for the Chromosomal Breaks in C-Myc That Lead to c-Myc/IgH Translocations’. Cell 135(6): 1028–38. doi:10.1016/j.cell.2008.09.062.

Roy, Bishnudeo, Swati Shukla, Marcin Łyszkiewicz, Martina Krey, Nuno Viegas, Sandra Düber, and Siegfried Weiss. 2009. ‘Somatic Hypermutation in Peritoneal B1b Cells’. Molecular Immunology 46(8–9): 1613–19. doi:10.1016/j.molimm.2009.02.026.

Rubtsova, Kira, Anatoly V. Rubtsov, Linda F. van Dyk, John W. Kappler, and Philippa Marrack. 2013. ‘T-Box Transcription Factor T-Bet, a Key Player in a Unique Type of B-Cell Activation Essential for Effective Viral Clearance’. Proceedings of the National Academy of Sciences 110(34): E3216–24. doi:10.1073/pnas.1312348110.

Shen, Qian, Hao Wang, Jonathan A. Roco, Xiangpeng Meng, Marita Bosticardo, Marie Hodges, Michael Battaglia, et al. 2025. ‘TCF1 and LEF1 Promote B-1a Cell Homeostasis and Regulatory Function’. Nature 646(8084): 442–51. doi:10.1038/s41586-025-09421-0.

Shinnakasu, Ryo, Takeshi Inoue, Kohei Kometani, Saya Moriyama, Yu Adachi, Manabu Nakayama, Yoshimasa Takahashi, et al. 2016. ‘Regulated Selection of Germinal-Center Cells into the Memory B Cell Compartment’. Nature Immunology 17(7): 861–69. doi:10.1038/ni.3460.

Song, Wenzhi, Olivia Q. Antao, Emily Condiff, Gina M. Sanchez, Irene Chernova, Krzysztof Zembrzuski, Holly Steach, et al. 2022. ‘Development of Tbet-and CD11c-Expressing B Cells in a Viral Infection Requires T Follicular Helper Cells Outside of Germinal Centers’. Immunity 55(2): 290–307.e5. doi:10.1016/j.immuni.2022.01.002.

Souza, Scott P., Samantha D. Splitt, Juan C. Sànchez-Arcila, Julia A. Alvarez, Jessica N. Wilson, Safuwra Wizzard, Zheng Luo, Nicole Baumgarth, and Kirk D. C. Jensen. 2021. ‘Genetic Mapping Reveals Nfkbid as a Central Regulator of Humoral Immunity to Toxoplasma Gondii’. PLoS Pathogens 17(12): e1010081. doi:10.1371/journal.ppat.1010081.

Srikakulapu, Prasad, Tanyaporn Pattarabanjird, Aditi Upadhye, Sai Vineela Bontha, Victoria Osinski, Melissa A. Marshall, James Garmey, et al. 2022. ‘B-1b Cells Have Unique Functional Traits Compared to B-1a Cells at Homeostasis and in Aged Hyperlipidemic Mice With Atherosclerosis’. Frontiers in Immunology 13. doi:10.3389/fimmu.2022.909475.

Stoeckius, Marlon, Christoph Hafemeister, William Stephenson, Brian Houck-Loomis, Pratip K. Chattopadhyay, Harold Swerdlow, Rahul Satija, and Peter Smibert. 2017. ‘Simultaneous Epitope and Transcriptome Measurement in Single Cells’. Nature Methods 14(9): 865–68. doi:10.1038/nmeth.4380.

Stone, Sara L., Jessica N. Peel, Christopher D. Scharer, Christopher A. Risley, Danielle A. Chisolm, Michael D. Schultz, Bingfei Yu, et al. 2019. ‘T-Bet Transcription Factor Promotes Antibody-Secreting Cell Differentiation by Limiting the Inflammatory Effects of IFN-γ on B Cells’. Immunity 50(5): 1172–1187.e7. doi:10.1016/j.immuni.2019.04.004.

Suan, Dan, Nike J. Kräutler, Jesper L. V. Maag, Danyal Butt, Katherine Bourne, Jana R. Hermes, Danielle T. Avery, et al. 2017. ‘CCR6 Defines Memory B Cell Precursors in Mouse and Human Germinal Centers, Revealing Light-Zone Location and Predominant Low Antigen Affinity’. Immunity 47(6): 1142–1153.e4. doi:10.1016/j.immuni.2017.11.022.

Takahashi, K, Y Kozono, T J Waldschmidt, D Berthiaume, R J Quigg, A Baron, and V M Holers. 1997. ‘Mouse Complement Receptors Type 1 (CR1;CD35) and Type 2 (CR2;CD21): Expression on Normal B Cell Subpopulations and Decreased Levels during the Development of Autoimmunity in MRL/Lpr Mice’. The Journal of Immunology 159(3): 1557–69. doi:10.4049/jimmunol.159.3.1557.

Talay, Oezcan, Donghong Yan, Hans D. Brightbill, Elizabeth E. M. Straney, Meijuan Zhou, Ena Ladi, Wyne P. Lee, et al. 2013. ‘Addendum: IgE+ Memory B Cells and Plasma Cells Generated through a Germinal-Center Pathway’. Nature Immunology 14(12): 1302–4. doi:10.1038/ni.2770.

Taylor, Justin J., Kathryn A. Pape, and Marc K. Jenkins. 2012. ‘A Germinal Center-Independent Pathway Generates Unswitched Memory B Cells Early in the Primary Response’. The Journal of Experimental Medicine 209(3): 597–606. doi:10.1084/jem.20111696.

Tedford, Kerry, Michael Steiner, Stanislav Koshutin, Karin Richter, Laura Tech, Yannik Eggers, Inga Jansing, et al. 2017. ‘The Opposing Forces of Shear Flow and Sphingosine-1-Phosphate Control Marginal Zone B Cell Shuttling’. Nature Communications 8(1): 2261. doi:10.1038/s41467-017-02482-4.

Tiller, Thomas, Makoto Tsuiji, Sergey Yurasov, Klara Velinzon, Michel C. Nussenzweig, and Hedda Wardemann. 2007. ‘Autoreactivity in Human IgG+ Memory B Cells’. Immunity 26(2): 205–13. doi:10.1016/j.immuni.2007.01.009.

Tirosh, Itay, Benjamin Izar, Sanjay M. Prakadan, Marc H. Wadsworth, Daniel Treacy, John J. Trombetta, Asaf Rotem, et al. 2016. ‘Dissecting the Multicellular Ecosystem of Metastatic Melanoma by Single-Cell RNA-Seq’. Science (New York, N.Y.) 352(6282): 189–96. doi:10.1126/science.aad0501.

Tomayko, Mary M., Natalie C. Steinel, Shannon M. Anderson, and Mark J. Shlomchik. 2010. ‘Cutting Edge: Hierarchy of Maturity of Murine Memory B Cell Subsets’. Journal of immunology (Baltimore, Md.: 1950) 185(12): 7146–50. doi:10.4049/jimmunol.1002163.

Tong, He, Li Wang, Kefan Zhang, Jing Shi, Yongshuai Wu, Yulong Bao, and Changshan Wang. 2023. ‘S100A6 Activates Kupffer Cells via the P-P38 and p-JNK Pathways to Induce Inflammation, Mononuclear/Macrophage Infiltration Sterile Liver Injury in Mice’. Inflammation 46(2): 534–54. doi:10.1007/s10753-022-01750-w.

Tornberg, U C, and D Holmberg. 1995. ‘B-1a, B-1b and B-2 B Cells Display Unique VHDJH Repertoires Formed at Different Stages of Ontogeny and under Different Selection Pressures.’ The EMBO Journal 14(8): 1680–89. doi:10.1002/j.1460-2075.1995.tb07157.x.

Vallejo, Jenifer, Ryosuke Saigusa, Rishab Gulati, Sujit Silas Armstrong Suthahar, Vasantika Suryawanshi, Ahmad Alimadadi, Christopher P. Durant, et al. 2022. ‘Combined Protein and Transcript Single-Cell RNA Sequencing in Human Peripheral Blood Mononuclear Cells’. BMC Biology 20: 193. doi:10.1186/s12915-022-01382-4.

Viant, Charlotte, Georg H. J. Weymar, Amelia Escolano, Spencer Chen, Harald Hartweger, Melissa Cipolla, Anna Gazumyan, and Michel C. Nussenzweig. 2020. ‘Antibody Affinity Shapes the Choice between Memory and Germinal Center B Cell Fates’. Cell 183(5): 1298–1311.e11. doi:10.1016/j.cell.2020.09.063.

Viant, Charlotte, Tobias Wirthmiller, Mohamed A. ElTanbouly, Spencer T. Chen, Ervin E. Kara, Melissa Cipolla, Victor Ramos, et al. 2021. ‘Germinal Center–Dependent and –Independent Memory B Cells Produced throughout the Immune Response’. Journal of Experimental Medicine 218(8): e20202489. doi:10.1084/jem.20202489.

Waffarn, Elizabeth E., Christine J. Hastey, Neha Dixit, Youn Soo Choi, Simon Cherry, Ulrich Kalinke, Scott I. Simon, and Nicole Baumgarth. 2015. ‘Infection-Induced Type I Interferons Activate CD11b on B-1 Cells for Subsequent Lymph Node Accumulation’. Nature Communications 6(1): 8991. doi:10.1038/ncomms9991.

Weisel, Florian J., Griselda V. Zuccarino-Catania, Maria Chikina, and Mark J. Shlomchik. 2016. ‘A Temporal Switch in the Germinal Center Determines Differential Output of Memory B and Plasma Cells’. Immunity 44(1): 116–30. doi:10.1016/j.immuni.2015.12.004.

Weisel, Florian, and Mark Shlomchik. 2017. ‘Memory B Cells of Mice and Humans’. Annual Review of Immunology 35(Volume 35, 2017): 255–84. doi:10.1146/annurev-immunol-041015-055531.

Weisel, Nadine M., Stephen M. Joachim, Shuchi Smita, Derrick Callahan, Rebecca A. Elsner, Laura J. Conter, Maria Chikina, et al. 2022. ‘Surface Phenotypes of Naive and Memory B Cells in Mouse and Human Tissues’. Nature Immunology 23(1): 135–45. doi:10.1038/s41590-021-01078-x.

Wong, Rachel, Julia A. Belk, Jennifer Govero, Jennifer L. Uhrlaub, Dakota Reinartz, Haiyan Zhao, John M. Errico, et al. 2020. ‘Affinity-Restricted Memory B Cells Dominate Recall Responses to Heterologous Flaviviruses’. Immunity 53(5): 1078–1094.e7. doi:10.1016/j.immuni.2020.09.001.

Zuccarino-Catania, Griselda V., Saheli Sadanand, Florian J. Weisel, Mary M. Tomayko, Hailong Meng, Steven H. Kleinstein, Kim L. Good-Jacobson, and Mark J. Shlomchik. 2014. ‘CD80 and PD-L2 Define Functionally Distinct Memory B Cell Subsets That Are Independent of Antibody Isotype’. Nature immunology 15(7): 631–37. doi:10.1038/ni.2914.

